# Auxin transport network underlies xylem bridge formation between the hemi-parasitic plant *Phtheirospermum japonicum* and host Arabidopsis

**DOI:** 10.1101/2019.12.26.889097

**Authors:** Takanori Wakatake, Satoko Yoshida, Ken Shirasu

## Abstract

Parasitic plants form vascular connections to host plants for efficient material transport. The haustorium is the responsible organ for host invasion and subsequent vascular connection. After invasion of host tissues, vascular meristem-like cells emerge in the central region of the haustorium, differentiate into tracheary elements, and establish a connection, known as a xylem bridge, between parasite and host xylem systems. Despite the importance of this parasitic connection, the regulatory mechanisms of xylem bridge formation are unknown. Here we show the role of auxin and auxin transporters during the process of xylem bridge formation using an Orobanchaceae hemiparasitic plant, *Phtheirospermum japonicum*. The auxin response marker DR5 has a similar expression pattern to tracheary element differentiation genes in haustoria. Auxin transport inhibitors alter tracheary element differentiation in haustoria, but biosynthesis inhibitors do not, demonstrating the importance of auxin transport during xylem bridge formation. The expression patterns and subcellular localization of PIN family auxin efflux carriers and AUX/LAX influx carriers correlate with DR5 expression patterns. The cooperative action of auxin transporters is therefore responsible for controlling xylem vessel connections between parasite and host.

## INTRODUCTION

Parasitic plants absorb water and nutrients from host plants by establishing an entry point to the host vascular system through a sophisticated invasive organ called a haustorium (Yoshida et al., 2016). The Orobanchaceae family includes various types of root parasitic plants such as obligate holoparasites that lack photosynthetic activity (e.g. *Orobanche spp.*), obligate hemiparasites (e.g. *Striga spp.*), and facultative hemiparasites (e.g. *Triphysaria* and *Phtheirospermum spp.*). These phanerogamic Orobanchaceae plants develop haustoria in their roots when in close proximity to host roots. Obligate parasites primarily form terminal haustoria at the tips of their radicles, but can also form secondary haustoria. The haustoria of facultative parasites and the secondary haustoria of obligate parasites emerge laterally from roots near the root apical meristem. Haustorium formation in Orobanchaceae parasites can be induced by chemical signals, collectively called haustorium inducing factors (HIFs), a group that includes quinones and phenolic acids (Goyet et al., 2019). Once parasite roots perceive HIFs, cell division and cell expansion are activated, and upon host attachment, apical haustorium cells differentiate into intrusive cells, which have characteristic elongated cell shapes, and function as the main cellular elements in host intrusion (Wakatake et al., 2018). A subset of intrusive cells that reaches host vasculature differentiates into conductive tracheary element xylem cells with thick secondary cell walls. Subsequently, haustorial cells located around the parasite root xylem and associated with the intrusive cells differentiate into tracheary elements and eventually connect to form a xylem bridge that establishes continuity between parasite and host xylem vessels (Ishida et al., 2016). Masses of tracheary elements called plate xylem also develop at the base of haustoria where the xylem bridge connects to the parasite root xylem (Dobbins and Kuijt, 1973).

The facultative parasite *Phtheirospermum japonicum* is an established model plant used to study the molecular mechanisms of plant parasitism and haustorium development (Cui et al., 2016; Ishida et al., 2011; Ishida et al., 2016; Spallek et al., 2017; Wakatake et al., 2018). *In vivo* imaging of cell cycle marker expression and nuclear tracking experiments indicate that multiple cell layers re-enter the cell cycle upon haustorium induction, and cell lineage analysis showed that haustorial epidermal cells differentiate into intrusive cells. After intrusive cells reach the host vasculature, cells in the central region of the haustorium and intrusive cells express procambium marker genes, such as *WOX4* and *HB15*, prior to xylem bridge formation (Wakatake et al., 2018). Some of these procambium-like cells further differentiate into tracheary elements accompanied by xylem cell marker gene expression, and establishment of the xylem bridge (Wakatake et al., 2018). Although some *Orobanche* spp. have a direct phloem connection with host plants (Aly et al., 2011; Zhou et al., 2004), *P. japonicum* does not form phloem-phloem connections, at least when parasitizing *Arabidopsis thaliana* (Spallek et al., 2017; Wakatake et al., 2018).

Auxin, or indol-3-acetic acid (IAA), regulates various developmental processes throughout the plant body (Teale et al., 2006). *In planta*, auxin concentration is tightly controlled by overlapping layers of regulation that affect rates of biosynthesis and catabolism, as well as intracellular and intercellular transport (Ruiz Rosquete et al., 2012; Zhao, 2018). Auxin transport is mainly mediated by the PIN-FORMED (PIN) proteins, a subset of the P-GLYCOPROTEIN (PGP) family, and by the AUXIN1/LIKE-AUX1 (AUX/LAX) family (Petrášek and Friml, 2009). PIN family proteins are classified depending on the length of the hydrophilic region flanked by two hydrophobic domains. ‘Long’ PIN subfamily members are localized in the plasma membrane and function as auxin efflux carriers, whereas ‘short’ PIN subfamily members are localized at the ER-membrane and are implicated in maintaining auxin homeostasis within a cell by intercellular auxin transport (Krecek et al., 2009). AUX/LAX proteins, which are similar to amino acid transporters, are localized at the plasma membrane and function as auxin influx carriers to regulate multiple aspects of plant development and tropism (Swarup and Péret, 2012). In contrast to these other transporters, the PGP class of ABC transporters is less well characterized. PGP1 and PGP19 export auxin, while PGP4 likely functions as a concentration-dependent bi-directional auxin transporter (Yang and Murphy, 2009).

Because of its ubiquity in development processes, it is not surprising that a link has been demonstrated between auxin and haustorium development in parasitic plants. Exogenous treatment with high concentrations of IAA, the auxin activity inhibitor *p*-chlorophenoxyisobutyric acid (PCIB), or the auxin transport inhibitor 1-*N*-naphthylphthalamic acid (NPA), dramatically reduced the frequency of infection of *A. thaliana* by *Orobanche aegyptiaca* (Bar-Nun et al., 2008). Similarly, in *Triphysaria versicolor,* PCIB or the auxin transport inhibitor 2,3,5-triiodobenzoic acid resulted in a reduction of haustorium formation induced by both host plants and the HIF 2,6-dimethoxy-ρ-benzoquinone (Tomilov et al., 2005). Auxin signaling and transport genes are upregulated in haustorial tissues of some parasitic plants (Ranjan et al., 2014; Zhang et al., 2015). In particular, a local auxin response maximum is newly established at the root surface facing the host root prior to haustorium development, and is accompanied by the epidermis-specific induction of the auxin biosynthesis enzyme *P. japonicum YUCCA 3* (*PjYUC3*) (Ishida et al., 2016). RNAi suppression of *PjYUC3* demonstrated that local auxin biosynthesis is an important step for haustorium initiation (Ishida et al., 2016). However, the roles of auxin in the establishment of the xylem bridge remain unknown. Here we investigated auxin response at the later stages of haustorium formation leading to xylem bridge formation (48∼72hpi), specifically by detailing the dynamics of auxin response related to xylem bridge formation and the role of auxin transporters during this process. This work provides the first molecular insights into tracheary element differentiation that lead to xylem bridge formation during plant parasitism.

## RESULTS

### Auxin response sites overlap with the tracheary element differentiation pattern

DR5 expression was detected as a pattern resembling that of xylem bridge formation (Fig. 1A-C). To confirm whether auxin-responding cells overlap with tracheary element differentiation patterns, we simultaneously observed the expression patterns of DR5 and *PjCESA7*, the latter of which encodes a cellulose synthase involved in secondary cell wall formation (Mitsuda et al., 2007; Taylor et al., 1999; Wakatake et al., 2018). *PjCESA7* expression was observed in cells that were also expressing *DR5* (Fig. 1D-F), suggesting that large increases in auxin lead to tracheary element differentiation leading to xylem bridge formation. The auxin biosynthesis gene *PjYUC3,* which is upregulated upon haustorium initiation (Ishida et al., 2016), was expressed only in the epidermis even at later stages of haustorium development (Fig. 1G-I). These observations suggest that auxin produced by *PjYUC3* alone does not account for the total auxin response during haustorium formation, especially in later stages, just before xylem bridge formation.

**Fig. 1.**
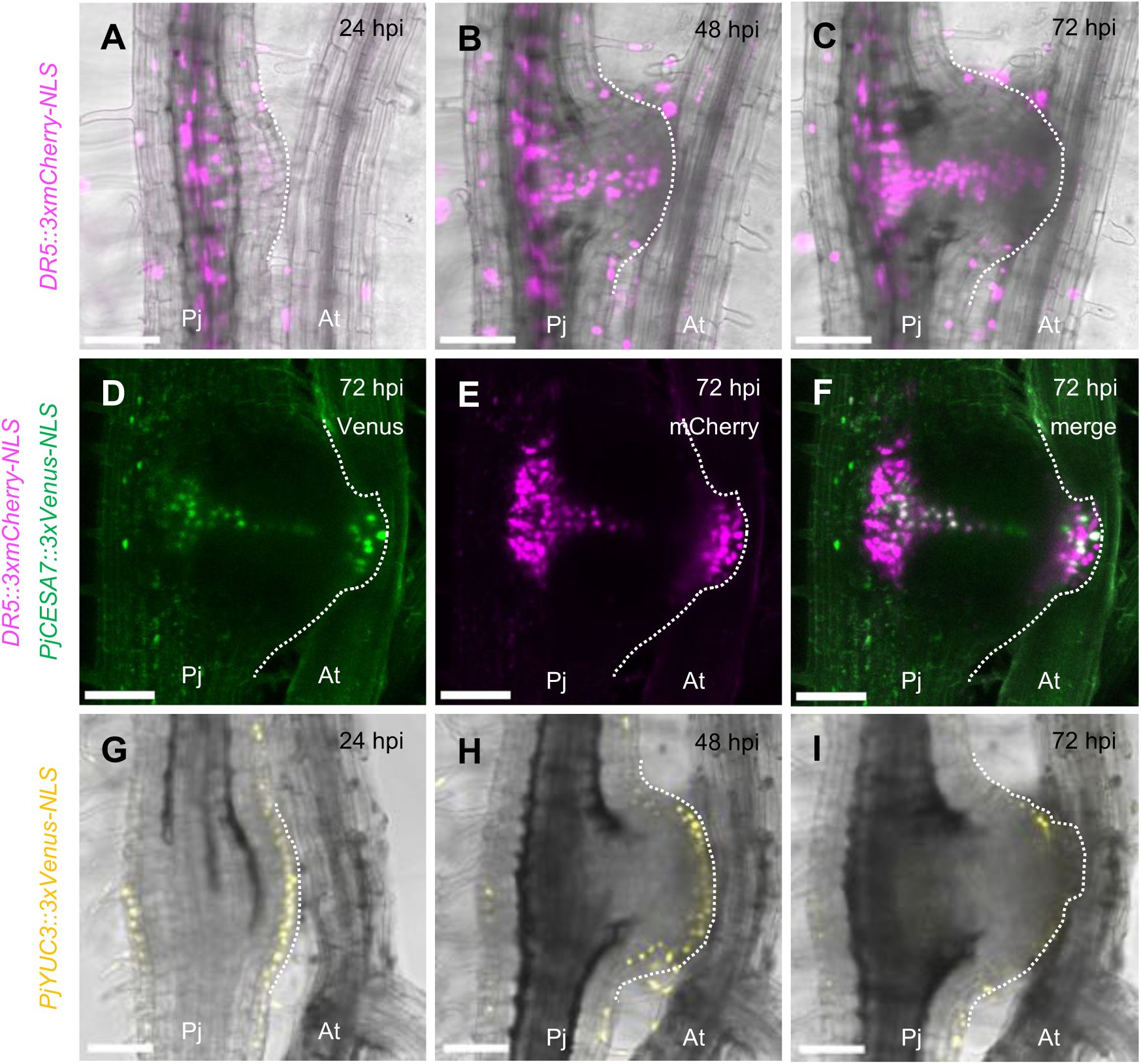
Auxin-responding cells respond to *PjCESA7* expression, but not to *PjYUC3*. (A)-(C) The expression pattern of *DR5::3xmCherry-NLS* during haustorium development from 24 to 72 hpi. A dotted line has been added to indicate the margin of the developing haustorium. (D)-(F) The expression pattern of *PjCESA7::3xVenus-NLS* and *DR5::3xmCherry-NLS* in the same haustorium at 72 hpi. (G)-(I) The expression pattern of *PjYUC3::3xVenus-NLS* during haustorium development from 24 to 72 hpi. Venus fluorescence is shown in green (D) and (F) or yellow (G)-(I) and mCherry fluorescence is indicated in magenta. Bright field images are merged with fluorescent images in (A)-(C) and (G)-(I). The same haustoria are shown in each time-course observation in (A)-(C) and (G)-(I), respectively. hpi, hours post infection; Pj, *P. japonicum* root; At, *A. thaliana* root. Bar = 100 µm.

### An auxin efflux inhibitor blocks xylem bridge formation

The auxin efflux inhibitor NPA and the influx inhibitor 3-chloro-4-hydroxyphenylacetic acid (CHPAA) were used to determine how auxin transport directionality affect haustorium formation (Katekar and Geissler, 1980; Parry et al., 2001) at 0, 24 and 48 hours post infection. Haustorial tissues were sampled at 96 hpi and stained with Safranin-O to visualize xylem bridge formation. NPA treatment of the haustorium at each time point effectively blocked the xylem bridge connection (Fig. 2A-C), although tracheary element differentiation was still observed in the intrusive region and in the plate xylem (Fig. 2D, E). Auxin transport within haustorium tissues is therefore likely important for xylem bridge formation, as NPA application above the haustorium on the parasite, or applied to the host did not block xylem bridge formation (Fig. S1A-C). Interestingly, CHPAA treatment did not block xylem bridge formation, but did affect its morphology (Fig. 2F). We also tested the effect of auxin biosynthesis inhibitors L-Kynurenine and Yucasin (He et al., 2011; Nishimura et al., 2014) on haustorium development. The combined treatment of two inhibitors at 0 hpi completely abolished haustorium initiation, whereas treatment at 48 hpi only partially blocked xylem bridge formation (Fig. S1D-F). Thus, we concluded that auxin biosynthesis inhibitors delay xylem bridge formation when applied at 48 hpi. This result demonstrates that *de novo* local auxin biosynthesis by epidermal cells is important for haustorium initiation as reported previously (Ishida et al., 2016), but is not essential for xylem bridge formation *per se*. Taken together, these inhibitor experiments suggest that auxin efflux activity within haustoria at later developmental stages (∼48 hpi) is crucial for xylem bridge formation.

**Fig. 2.**
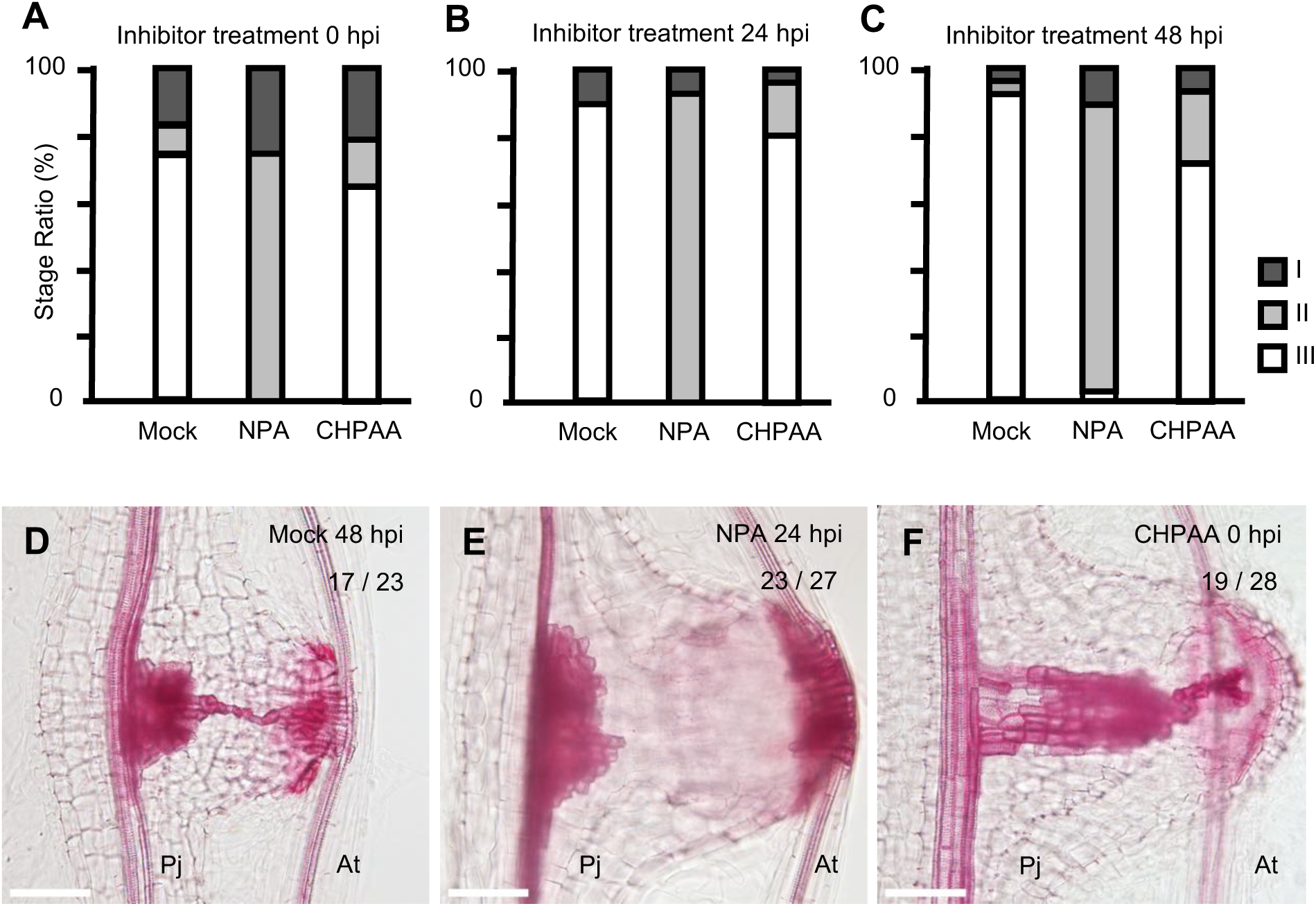
The effect of auxin transport inhibitors on haustorium development. (A)-(C) Ratio of each developmental stage of xylem bridge formation at 96 hpi after auxin transport inhibitors treatments (I, pre-initiation; II, developing; III, fully connected). Treatment concentrations are 5 µM for NPA and 10 µM for CHPAA. Chemical treatments were started at 0 hpi (A), 24 hpi (B), or 48 hpi (C). n = 23 ∼ 30 for each chemical treatment. (D)-(F) Representative photos of xylem bridge formation for mock (D), NPA (E), and CHPAA (F) treatment. The fraction of samples showing a similar pattern is shown in the image. Tracheary elements were stained red with Safranin-O. hpi, hours post infection; Pj, *P. japonicum* root; At, *A. thaliana* root. Bar = 100 µm.

### Expression and localization of PIN proteins during haustorium development

To identify which auxin efflux carriers function during xylem bridge formation, we measured the expression of a family of auxin transporters encoded by the *PIN* genes, which direct auxin flow in plants (Petrášek et al., 2006; Wisniewska et al., 2006). Six *PIN* genes encoding canonical domain structures (two hydrophobic regions flanking an intercellular hydrophilic loop) were identified in the *P. japonicum* draft genome (Fig. S2) (Conn et al., 2015; Krecek et al., 2009). All six *PjPIN* genes encode the ‘long’ subclade of *PINs*. PjPIN6 was omitted from this study because its ortholog, AtPIN6, localizes to both the plasma membrane and the ER-membrane, and it has a different function among the PIN family (Simon et al., 2016). The expression patterns and subcellular localizations of the five PIN proteins (PjPIN1 to PjPIN4, and PjPIN9) were monitored during haustorium formation by fusing the Venus protein into the middle of a hydrophilic loop, similar to what was previously reported (Benková et al., 2003; Blilou et al., 2005; Vieten et al., 2005; Xu and Scheres, 2005; Žádníková et al., 2010).

Without a host, PjPIN1 expression was detected ubiquitously in the root tip, except in the epidermis, and was strongest in stele tissues around the elongation zone (Fig. 3A). PjPIN1-Venus localized at the rootward plasma membrane in stele tissues, similar to AtPIN1 (Gälweiler et al., 1998). By contrast, *PjPIN2* expression was detected only in epidermis and the adjacent cortex layer in *P. japonicum* roots when grown without a host (Fig. 3B). PjPIN2-Venus is localized within the shootward membrane in epidermis, and at the lateral side of the membrane specifically in the transition zone, which is similar to localization of the AtPIN2 protein (Fig. 3B) (Müller et al., 1998). PjPIN9, a close homolog of tomato SIPIN9 or ‘Sister-of-PIN1a’ (SoPIN1a) (Solyc10g078370) (O’Connor et al., 2014), has no ortholog in Arabidopsis (Fig. S2). It was expressed in the meristematic zone of vascular tissues, and was localized at the rootward side of the membrane grown without a host (Fig. 3C).

**Fig. 3.**
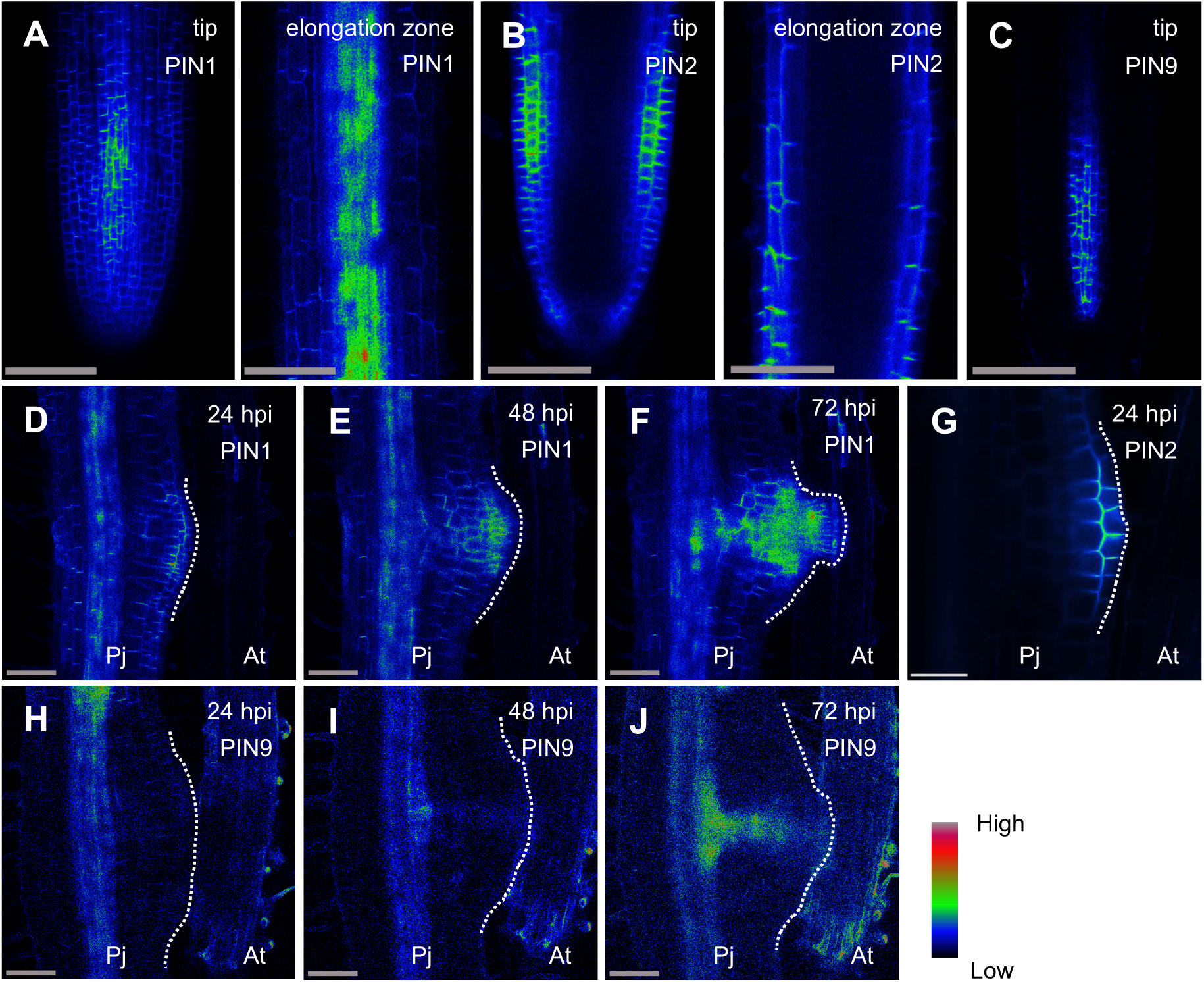
Expression patterns of PjPIN1, PjPIN2, and PjPIN9 in *P*. *japonicum* roots and haustoria. (A) and (D)-(F) *PjPIN1::PjPIN1-Venus* expression in *P. japonicum* root tips and elongation zones (A), and during haustorium formation at 24, 48 or 72 hpi (D)-(F). (B) and (G) *PjPIN2::PjPIN2-Venus* expression in *P. japonicum* root tips and elongation zones (B), and in haustoria at 24 hpi (G). (C) and (H)-(J) *PjPIN9::PjPIN9-Venus* expression in *P. japonicum* root tips (C) and during haustorium formation at 24, 48 or 72 hpi (H)-(J). Venus fluorescence intensity is depicted in a Rainbow RGB spectrum. The white dotted lines outline haustoria. hpi, hours post infection; Pj, *P. japonicum* root; At, *A. thaliana* root. Bar = 100 µm.

Upon infection, very weak PjPIN1 expression in the cortex of the haustorium was detected at 24 hpi (Fig. 3D), but expression became stronger in the central region toward the host side at 48 and 72 hpi (Fig. 3E,F). In the case of PjPIN2, fluorescence at 24 hpi was localized on the rootward side of epidermal cells above the haustorium apex, but on the shootward side of cells below the haustorium apex (Fig. 3G). PjPIN2 was also localized at the inner and outer lateral sides of the epidermal membrane and the neighboring cortical layer of the haustorium apex, respectively. PjPIN2 expression could not be detected in haustorium tissues at later stages. PjPIN9 expression was observed in stele tissues, but not in haustoria during the early stages of development (24 hpi and 48 hpi) (Fig. 3H, I). At later stages of haustorium development (72 hpi), PjPIN9 fluorescence was detected in the plate xylem toward the center of the haustorium (Fig. 3J). These expression analyses indicate that PjPIN2 plays a critical developmental role at the haustorial apex at an early stage, but PjPIN1 and PjPIN9 take over responsibility for auxin transport during later stages of development. For PjPIN3 and PjPIN4, we did not detect significant changes in haustorium expression patterns, suggesting that these transport proteins are not involved in haustorium formation (Fig. S3).

To gain further insight into the timing and extent of auxin flow directed by PjPIN1 and PjPIN9 prior to xylem bridge formation in the haustorium, we sectioned haustoria to observe the subcellular localization of PjPIN1-Venus and PjPIN9-Venus (Fig. 4, full stack images are shown in Fig. S4 and S5). Haustorial cells are rather unorganized and pleiomorphic. Nevertheless, we found polar localizations of PjPIN1 and PjPIN9. PjPIN1-Venus was localized at the inner lateral side of the plasma membrane in haustorial cells near host roots (Fig. 4A) and at the lateral membrane of parasite pericycle cells, proximal to the area where the plate xylem develops (Fig. 4B). In contrast, PjPIN9-Venus was detected at the lateral membrane facing the center of haustoria in cells at the edge of the procambium-like domain (Fig. 4C), and at the lateral membrane proximal to the area where the plate xylem develops near the parasite vasculature (Fig. 4D). These PjPIN1 and PjPIN9 localizations correspond to tracheary element differentiation sites, corroborating the idea that these two proteins function in xylem bridge formation during infection.

**Fig. 4.**
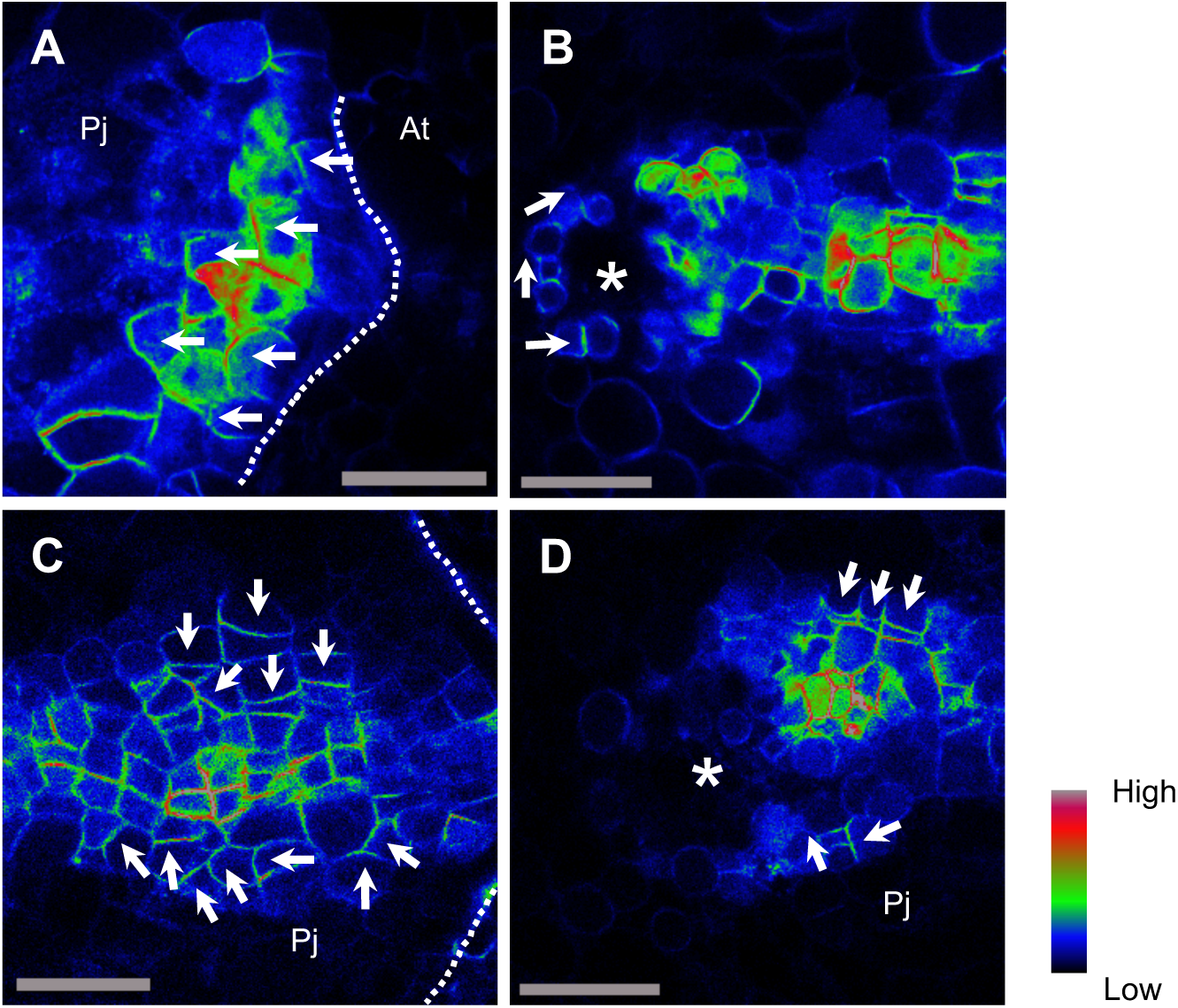
Subcellular localization of PjPIN1 and PjPIN9 in haustoria before xylem bridge formation. (A) and (B) *PjPIN1::PjPIN1-Venus* expression in cross section of the haustorium near host tissue (A), and near parasite vasculature (B). (C) and (D) *PjPIN9::PjPIN9-Venus* expression in cross section of the haustorium near host tissue (C), and near the parasite vasculature (D). Full stack images are in supplementary figures 4 and 5. Venus fluorescence intensity is depicted in a Rainbow RGB spectrum. Arrows indicate polar localization of PIN-Venus protein fusions. Asterisks indicate parasite root vasculature. The white dotted lines mark the haustorial interface with host tissue. Pj, *P. japonicum* root; At, *A. thaliana* root. Bar = 50 µm.

**Fig. 5.**
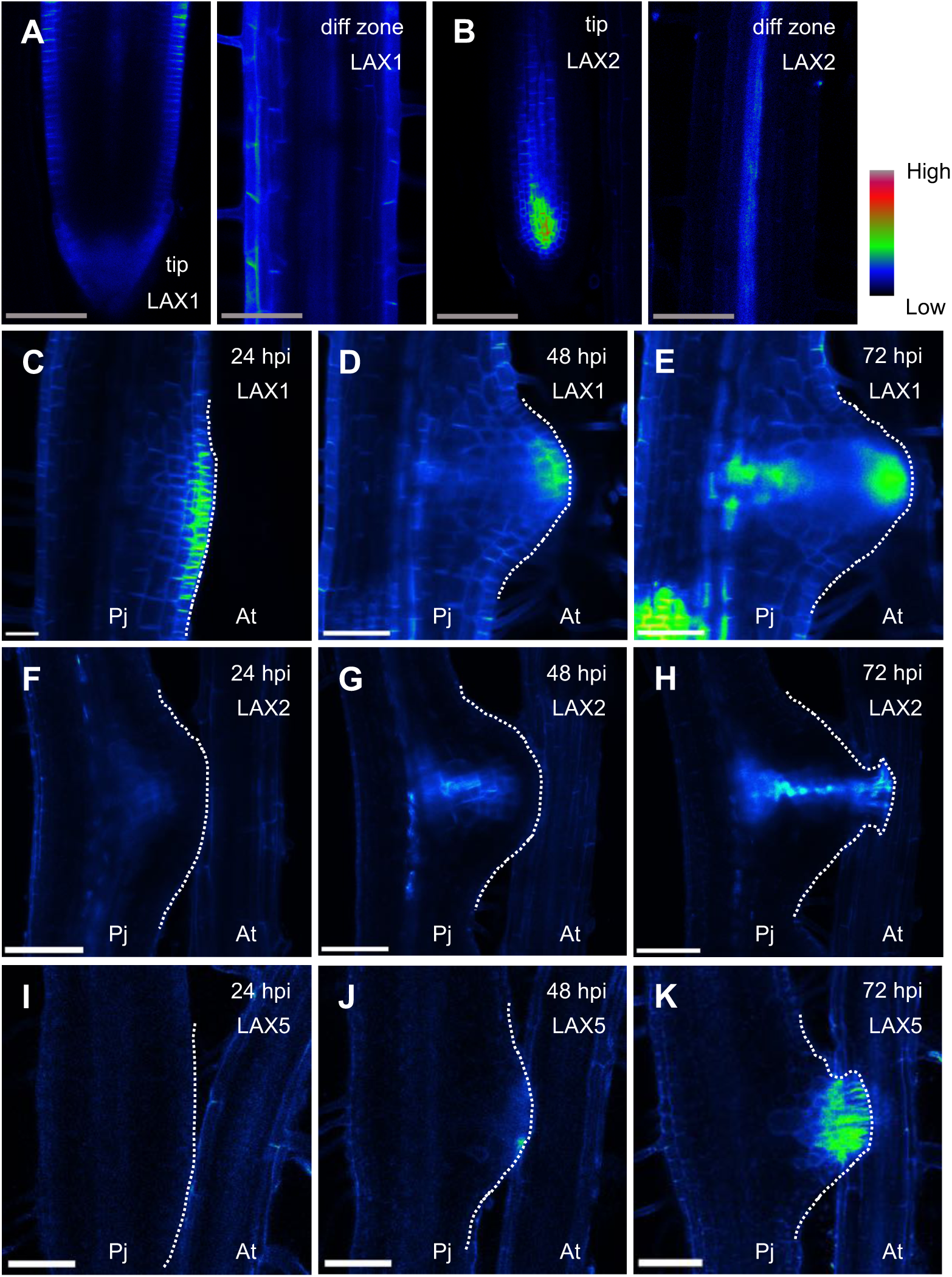
Expression patterns of PjLAX1, PjLAX2, and PjLAX5 in *P*. *japonicum* roots and haustoria. (A) and (C)-(E) *PjLAX1::PjLAX1-Venus* expression in *P. japonicum* root tip and differentiation zone (A), and during haustorium formation at 24, 48 or 72 hpi (C)-(E). (B) and (F)-(H) *PjLAX2::PjLAX2-Venus* expression in *P. japonicum* root tip and differentiation zone (B), and during haustorium formation at 24, 48 or 72 hpi (F)-(H). (I)-(K) *PjLAX5::PjLAX5-Venus* expression during haustorium formation at 24, 48 or 72 hpi. White dotted lines mark the outline of haustoria. Venus fluorescence intensity is depicted in a Rainbow RGB spectrum. hpi, hours post infection; Pj, *P. japonicum* root; At, *A. thaliana* root. Bar = 100 µm.

### Expression of the LAX gene family during haustorium development

Because other auxin influx transporters may be involved in regulating haustorium development, we investigated the expression patterns of the *AUX/LAX* gene family (hereafter denoted as *LAX*), a group that encodes auxin influx transporters (Carrier et al., 2008; Swarup et al., 2008; Yang et al., 2006). Five LAX genes were identified in the *P. japonicum* draft genome (Fig. S6) (Conn et al., 2015; Yoshida et al., 2019; Young et al., 1999). The expression patterns and subcellular localization of each LAX protein was monitored during haustorium formation by fusing Venus in a similar position to previously reported fluorescent protein-tagged LAX proteins in *A. thaliana* (Péret et al., 2012; Swarup et al., 2004; Swarup et al., 2008). The expression patterns of PjLAX1, PjLAX2, and PjLAX5 (Fig. 5), but not PjLAX3 or PjLAX4 (Fig. S7), were significantly different in haustorium formation. Without a host, PjLAX1 was expressed in epidermis, root cap, and pericycle in *P. japonicum* roots (Fig. 5A). With a host, strong PjLAX1 expression was observed at the haustorium apex (Fig. 5C-E). At later developmental stages (72 hpi), weak fluorescent signals were also observed in the central region of the haustorium (Fig. 5E). For PjLAX2, strong expression was observed in vascular tissues above the quiescent center (QC), and weak expression was observed above the meristematic zone when no host was present (Fig. 5B). With a host, *PjLAX2* expression gradually expanded from the parasite root vasculature tissues (24 hpi, Fig. 5F) to the central haustorium region (48 hpi, Fig. 5G), and eventually to the intrusive cells (72 hpi, Fig. 5H). *PjLAX2* expression before xylem bridge formation was very similar to that of DR5 (Fig. 1, 5H). In contrast, PjLAX5 expression was not detected in *P. japonicum* roots before haustorial induction, nor during the early stages of haustorium development (24 hpi, Fig. 5I). At around 48 hpi, when the haustorium begins host tissue invasion, weak expression was observed at the interface with host tissues (Fig. 5J). Strong *PjLAX5* expression was detected in intrusive cells as they developed (72 hpi, Fig. 5K). From these expression analyses, we concluded that PjLAX1, PjLAX2, and PjLAX5 likely fine-tune the auxin concentration gradient to initiate xylem bridge connections.

### The effect of auxin transport inhibitors on reporter expression

As a way to test our hypothesis that auxin transport is the main driver of the timing and direction of vascular bridge formation, we used auxin transport inhibitors to alter auxin responses in haustoria by treating *P. japonicum* roots expressing the DR5 reporter with NPA and CHPAA during haustorium development. DR5 expression near the parasite root vasculature was not affected by NPA treatment. However, no Venus signal was observed in the central region of haustoria (Fig. 6A-C). In contrast, CHPAA treatment did not disturb DR5 expression in the central region of the haustorium (Fig. 6D-E). Furthermore, normal xylem bridge formation was confirmed with CHPAA treatment but not with NPA treatment at 96 hpi (Fig. 6C,F), as seen in non-transgenic *P. japonicum* (Fig. 2E-F).

**Fig. 6.**
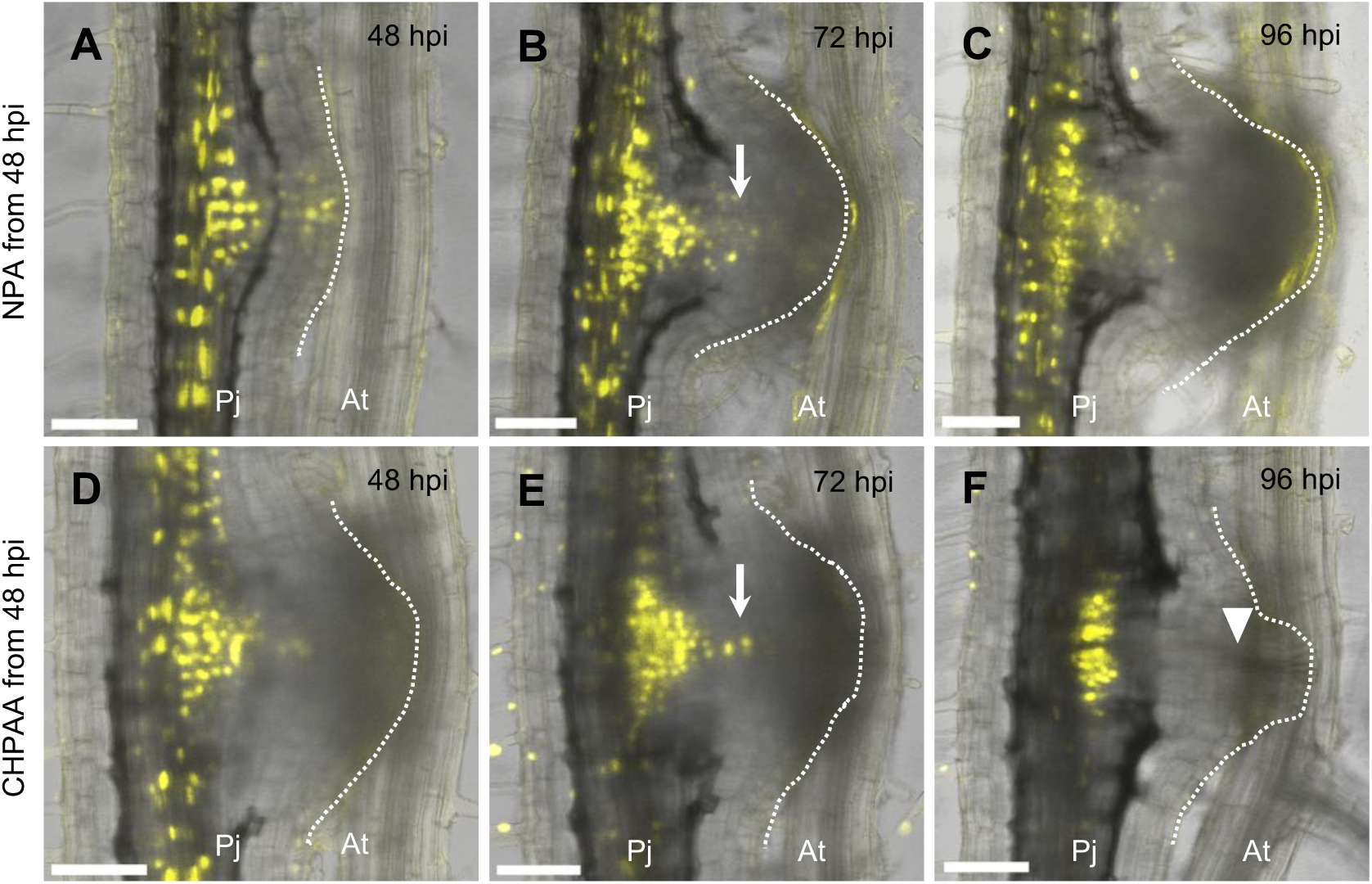
*DR5::3xVenus-NLS* expression during haustorium development with auxin inhibitor treatments. Expression patterns of *DR5::3xVenus-NLS* during haustorium development with exogenous application of 5 µM NPA at 48 hpi (A) to (C) or 10 µM CHPAA at 48 hpi (D) to (F) at 24, 48 or 72 hpi. Arrows indicate DR5 expression in the haustorial central region. An arrowhead indicates the nascent xylem bridge. White dotted lines mark the outline of the developing haustoria. Venus fluorescence is in yellow. Bright field images and Venus fluorescent images are merged. hpi, hours post infection; Pj, *P. japonicum* root; At, *A. thaliana* root. Bar = 100 µm.

NPA treatment not only blocked xylem bridge formation, but also made haustoria larger when treated from the beginning of the experiment (i.e. at 0 hpi) (Fig. 2E). We further assessed the effects of auxin transport inhibitors on haustorium tissue structure by employing *PjHB15a* as a marker for procambium-like cells (Wakatake et al., 2018). Although the temporal expression patterns of *PjHB15a* were not altered by NPA treatment, the signal was more diffuse in haustoria than in untreated plants (Fig. 7A-G). *PjHB15a* expression was weaker at the plate xylem development site when treated with CHPAA (Fig. 7H-K). Cross sections of a haustorium also revealed that *PjHB15a*-positive procambium-like cells occupied a much larger area of tissues with NPA than with CHPAA treatment (Fig. 7L,M), resulting in larger haustoria. These results indicate that even in the presence of auxin transport inhibitors, procambium-like cells are able to form in the haustorium.

**Fig. 7.**
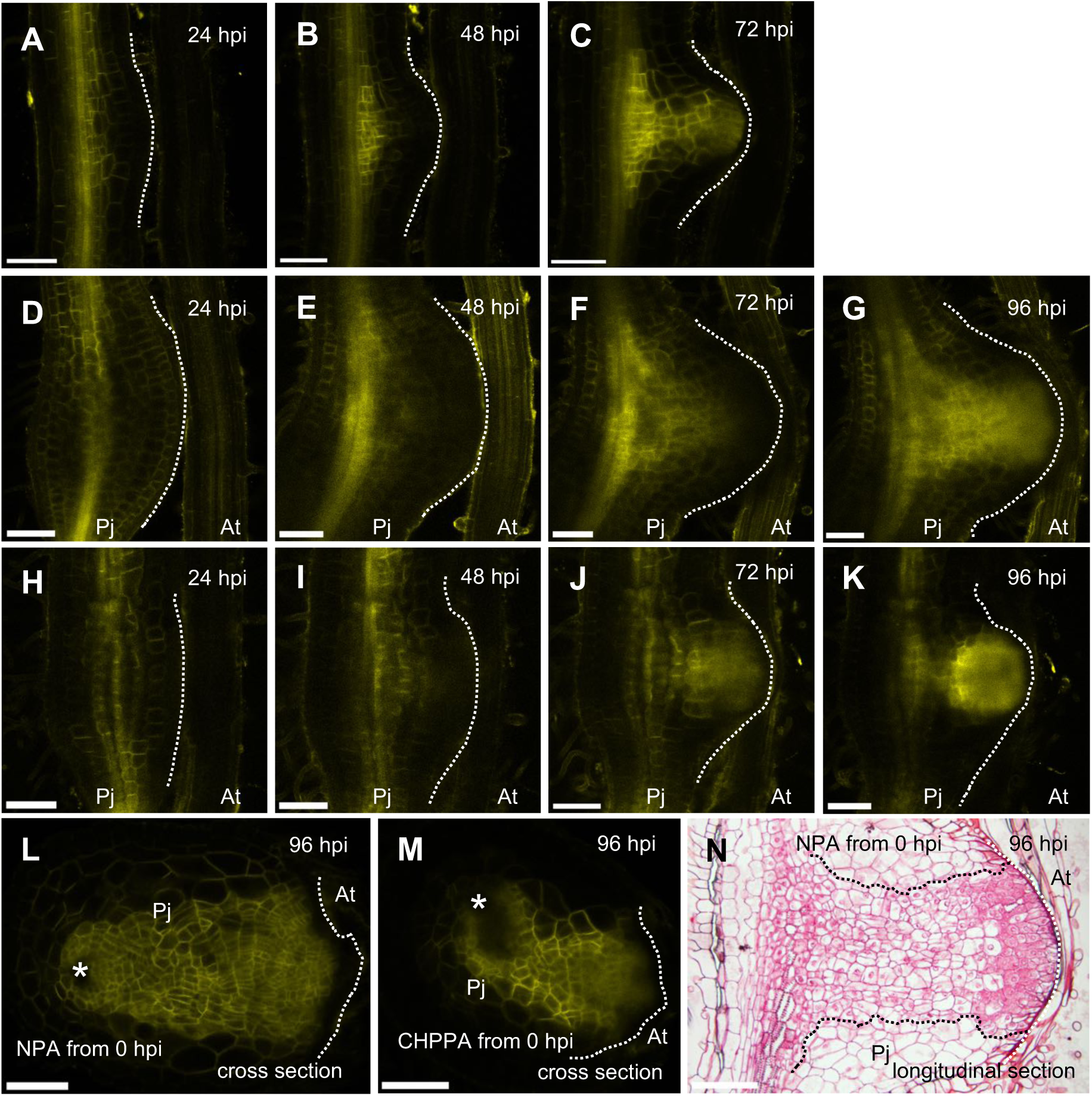
*PjHB15a::3xVenus-SYP* expression during haustorium development with auxin inhibitor treatments. Expression pattern of *PjHB15a::3xVenus-SYP* during haustorium development at 24, 48 or 72 hpi following mock chemical treatment at 0 hpi (A) to (C), 5 µM NPA at 0 hpi (D) to (G) and (L), or 10 µM CHPAA at 0 hpi (H) to (K) and (M). Cross sections of haustoria are shown in (L) and (M). Asterisks indicate parasite root vasculature. Venus fluorescence is shown in yellow. (N) Longitudinal section of a haustorium at 96 hpi treated with 5 µM NPA at 0 hpi. The section was stained with Safranin-O. Black dotted lines surround procambium-like cells in the haustorium. White dotted lines mark the outline of haustoria. hpi, hours post infection; Pj, *P. japonicum*; At, *A. thaliana* root. Bar = 100 µm.

### *PjPIN1* and *PjPIN9* are involved in xylem bridge formation

Based on our expression analyses, and confirmed by the inhibitor experiments, *PjPIN1* and *PjPIN9* are likely required for xylem bridge formation. To assess the function of *PjPIN1* and *PjPIN9* during haustorium development, we knocked down these genes using RNAi with the pHG8-YFP vector (Bandaranayake et al., 2010). Highly variable sequences located in the hydrophilic loop of the *PIN* genes were cloned into the pHG8-YFP vector to form pHG8-PIN1 and pHG8-PIN9 as a way to avoid possible off-target effects. We confirmed reduced expression of the targeted genes in hairy roots with RT-qPCR (Fig S8A,B). Hairy roots harboring pHG8-PIN1 and pHG8-PIN9 both had fewer successful xylem connections at 96 hpi (52.7% and 59.6%, respectively) compared with the empty vector control (92.3%) (Fig. 8A-D). Notably, we observed partial tracheary element differentiation in the plate xylem in *PIN1*-knockdown roots (Fig. 8C-II). In contrast, partial tracheary element differentiation was observed only in the intrusive region of *PIN9*-knockdown roots (Fig. 8D-II). Importantly, several connected xylem bridges lacked plate xylems in *PIN9*-knockdown roots (Fig. 8D-III). Taken together, these data indicate that both *PjPIN1* and *PjPIN9* positively regulate xylem bridge formation, but play different roles in haustorium development.

**Fig. 8.**
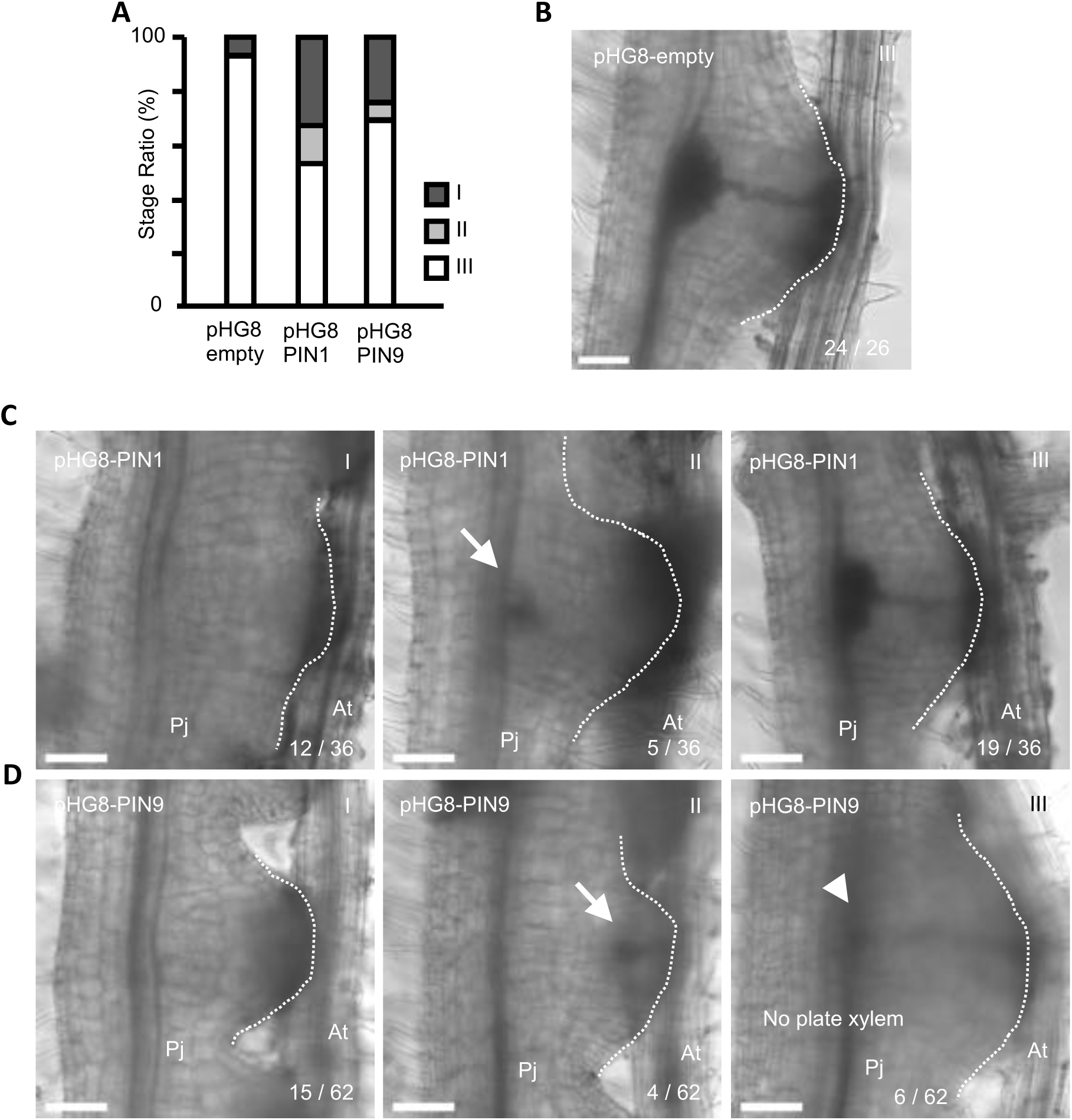
PjPIN1 and PjPIN9 are involved in xylem bridge formation. (A) Ratio of each developmental stage of xylem bridge formation in hairy roots at 96 hpi (I, pre-initiation; II, developing; III, fully connected). n = 26∼62. (B)-(D) Representative photos of each developmental stage in hairy roots transformed with pHG8-empty vector (B), pHG8-PIN1 (C), and pHG8-PIN9 (E). Arrows indicate partial differentiation of tracheary elements. An arrowhead points to where the plate xylem should have formed. White dotted lines mark the outline of haustoria. Pj, *P. japonicum*; At, *A. thaliana* root. Bar = 100 µm.

## DISCUSSION

### Formation of two auxin maxima in haustoria

Our previous work showed that locally induced auxin biosynthesis is an important step for haustorium initiation in parasitic plants (Ishida et al., 2016). In the present study, we investigated the role of auxin in later stages of haustorium development and found that auxin flux coordinated by auxin transporters is crucial for xylem bridge formation between a parasitic plant and its host. One peculiar phenomenon during haustorium development is the formation of two distinct auxin maxima, one at the haustorium apex where intrusive cells face host tissues, and the other at the site where the plate xylem forms adjacent to parasite root vasculature (Fig. 1E). High numbers of tracheary elements are differentiated at these two sites (Fig. 2D). The auxin efflux transport inhibitor NPA does not block formation of tracheary elements at the two sites, but instead increases the area composed of tracheary elements at each site (Fig. 2E), suggesting that a key function of auxin efflux transporters in the haustorium is to sharpen the area of auxin maxima at these sites, especially during the early stages of haustorium development. The auxin efflux transporters PjPIN1 and PjPIN2 were consistently expressed in haustorial apices at early stages (24 hpi, Fig. 3D,G) and PjPIN9 was expressed around the plate xylem development site at later stages (48∼ hpi, Fig. 3I,J). Importantly, PjPIN2 localized polarly toward the haustorium apex in the epidermis, but in the outer cortex layer it localized only at the outer lateral side (Fig. 3G). Similarly, PjPIN9 localized at the lateral membrane facing the plate xylem area in haustorial cells near the parasite root vasculature (Fig. 4D). Thus, PjPIN2 and PjPIN9 likely contribute to auxin concentrations at the two auxin maximum sites. Particularly in *PjPIN9*-knockdown roots, we observed the absence of the plate xylem (Fig. 8D-III), further supporting the idea that PjPIN9 is required for establishing higher auxin concentrations at the plate xylem development site.

LAX auxin influx transporters are also likely involved in tracheary element patterning at the haustorium apex and at the plate xylem. CHPAA treatment disturbs typical xylem bridge patterning in these areas (Fig. 2F). PjLAX1 was highly expressed at the haustorium apex and the plate xylem formation site (Fig. 5C-E). PjLAX2 was also strongly expressed at the plate xylem development site (Fig.5F-H). Unlike PjLAX1 and PjLAX2, which were also expressed in normal roots without a host, PjLAX5 exhibited a haustorium-apex-specific expression pattern (Fig. 5K). These expression patterns are consistent with DR5 auxin reporter patterns in haustoria (Fig. 1E), indicating that they contribute to the maintenance of higher auxin concentrations at these sites.

### Xylem bridge formation

The most conspicuous effect of NPA treatment on haustorium development is the failure of xylem bridge connections between the two auxin maxima sites (Fig. 2E). By contrast, CHPAA treatment did not block xylem bridge formation, but instead broadened the xylem bridge (Fig. 2F). Thus, auxin efflux activity is essential for connecting the two auxin maxima by the xylem bridge, whereas influx activity is important for restricting the bridge formation area, resulting in a single cell-wide xylem bridge in the region of the haustorium closest to the parasite. PjPIN1 and PjPIN9 were expressed in the middle of the haustorium prior to formation of the xylem bridge, therefore these proteins likely act during xylem bridge formation (Fig. 3D-, H-J). PjPIN1 is preferentially expressed near the host plant (Fig. 3F), whereas PjPIN9 is expressed more strongly near the parasite vasculature (Fig. 3J), suggesting that these two PIN proteins play different roles in maintaining the auxin gradient in haustorial tissues. PjPIN1 shows subcellular localization toward the parasite vasculature in the areas nearest host tissues, suggesting an inward auxin flow from the haustorium apex (Fig. 4A,B). PjPIN9 localizes within the membrane facing toward the xylem bridge formation site, indicating that there is additional auxin flow toward the haustorium center (Fig. 4C,D). Indeed, knockdown of each gene leads to two distinct phenotypes that are consistent with their expression patterns and subcellular localizations. In *PjPIN1*-knockdown roots, a partial xylem bridge only formed in the plate xylem (Fig. 8C-II). In contrast, a partial xylem bridge formed only in the intrusive cell region in *PjPIN9*-knockdown roots (Fig. 8D-II), and some successfully connected xylem bridges lacked the plate xylem altogether (Fig. 8D-III). These observations suggest that PjPIN1 and PjPIN9 coordinately function to direct auxin flow between parasite vasculature and host vasculature. Some caution should be taken when interpreting the NPA treatment results, as NPA inhibits IAA export by PGP proteins as well as by PIN proteins (Bouchard et al., 2006; Geisler et al., 2005; Petrášek et al., 2006; Yang and Murphy, 2009). However, unlike PINs, PGP proteins are evenly distributed throughout the plasma membrane and are therefore less likely to contribute to shaping auxin gradients (Cho and Cho, 2013). Still, the possibility that the effect of NPA on xylem bridge formation is due to an inhibitory effect on PGP proteins in addition to PjPIN1 and PjPIN9 cannot be excluded from the experiments described here.

### A model for auxin transport networks in haustoria

Based on these experimental results, we propose a model to describe how the auxin transport network distributes auxin to form the xylem bridge during haustorium development (Fig. 9A-D). In early developmental stages, host-inducible PjYUC3 produces auxin locally in the epidermis, and PjPIN2 together with PjLAX1 maintain high auxin concentrations at the haustorium apex (Fig. 9A). As the haustorium develops, PjPIN1 directs auxin flow from its apex toward the parasite’s main vasculature, while PjPIN9 further restricts auxin movement to create a higher auxin gradient at the plate xylem formation site (Fig. 9B). At the same time, PjLAX2 expression expands from the parasite vasculature toward the haustorium apex to maintain a high auxin gradient in the central region. At later stages, PjLAX1 and PjLAX5 maintain high auxin concentrations at the haustorium apex. The xylem bridge is coordinately formed with the auxin gradient shaped by these auxin carriers (Fig. 9C). With NPA treatment, high auxin response sites are not established in the central region and thus procambium-like cells cannot further differentiate into tracheary elements (Fig. 9D).

**Fig. 9.**
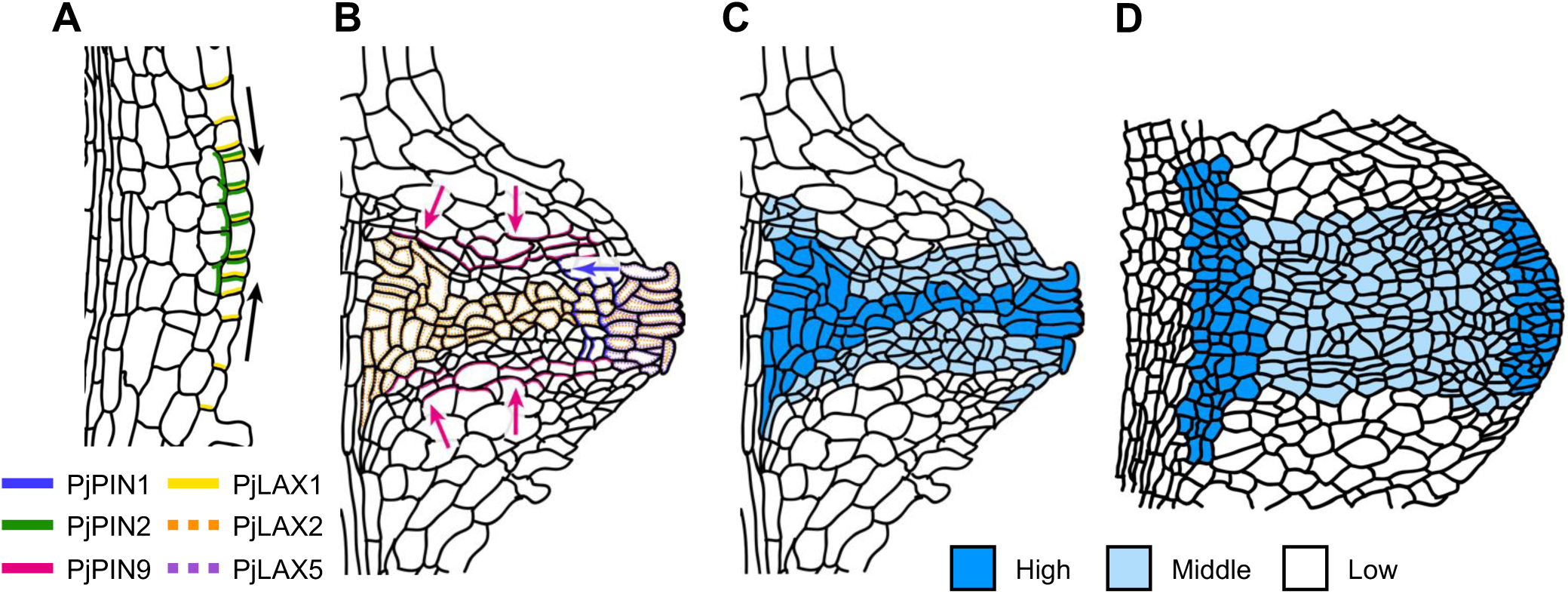
A model of auxin flow and concentration gradient during haustorium formation. (A) and (B) Schematic representation of subcellular localization and expression pattern of auxin transporters during haustorium development during early (A) and later stages (B). Lines indicate the subcellular localization of color-coded transporters as shown in the key. Dashed lines indicate the location of *PjLAX2* and *PjLAX5* gene expression. Arrows indicate the direction of auxin flow coordinated by PjPIN2 and PjLAX1 (black), PjPIN1 (blue), and PjPIN9 (magenta). (C) and (D) Schematic representation of the auxin concentration gradient in haustoria prior to xylem bridge formation (C) and in NPA-treated haustoria (D). The color scale indicates auxin levels.

### How did the haustorial auxin transportation system evolve?

The observation that PjPIN1, PjPIN2, PjPIN9, PjLAX1, PjLAX2 and PjLAX5 are expressed in haustoria, an organ that is formed only in parasitic plants, raises the question of how these proteins could have acquired these specific expression patterns. It is possible that the haustorium-specific transcriptional regulatory system of PIN and LAX genes was evolved independently within the Orobanchaceae lineage. However, PjPIN1 and PjPIN2 had similar expression and subcellular localization patterns around the root tip in the absence of a host as the Arabidopsis homologs AtPIN1 and AtPIN2 (Fig. 3A,B) (Blilou et al., 2005; Omelyanchuk et al., 2016; Vieten et al., 2005). Similarly, PjLAX1 and PjLAX2 are expressed much as their corresponding orthologs are in Arabidopsis when no host is present (Fig. 5A,B) (Péret et al., 2012; Swarup et al., 2001). Therefore, the regulatory *cis*-element and transcriptional factors controlling expression of these genes should be conserved in non-parasitic and parasitic plants. In the case of PjPIN9, there is no apparent ortholog in Arabidopsis, but it is closely related to the SoPIN1 clade that was lost from the Brassicaceae lineage (Fig. S2) (O’Connor et al., 2014). The role of SoPIN1 in roots is as yet unknown. Without a host, PjPIN9 was strongly localized to the rootward side in vascular tissues just above QC, suggesting that PjPIN9 likely directs auxin flow rootward to maintain a high auxin concentration in the root apical meristem (Fig. 3C). Thus, it is difficult to conclude whether PjPIN9 evolved specifically for parasitism. Similarly, while PjLAX5 does not have a clear ortholog in Arabidopsis, it does share a clade with some Fabaceae and *Populus trichocarpa* LAX genes (Fig. S6), indicating that the PjLAX5 lineage is not specific to Orobanchaceae parasitic plants.

Another possibility is that these proteins are not under haustorium-specific transcriptional or post translational regulation. Instead, these proteins may just follow the canalization pattern proposed by Sachs (Bennett et al., 2014; Sachs, 1969). In the Sachs model, auxin ‘sources’ (regions of high auxin concentration or production) can be connected to ‘sinks’ (regions of low auxin concentration or high turnover) in a self-organizing manner. In normal systems, developing tissues function as an auxin source and established vasculature functions as a sink. In parasitic plants, the haustorium apex may act as a source and the plate xylem formation site near root vasculature as a sink. In this scenario, auxin transporters could self-organize to form a xylem bridge between the haustorium apex as a source, and the plate xylem as a sink. Further studies of auxin transporters in other Orobanchaceae plants may clarify the origin of auxin transport systems in the haustoria of parasitic plants.

## MATERIALS AND METHODS

### Chemicals and reagents

All reagents were purchased from Wako Pure Chemical Industries, Ltd. (Osaka, Japan) unless otherwise stated.

### Plant materials and growth conditions

*P. japonicum* seeds were surface sterilized with 10% commercial bleach (Kao, Japan) then rinsed five times with sterile water, and sown directly on half-strength Murashige-Skoog (MS) agar plates (0.8% agar and 1% sucrose) for transformation experiments, and on nylon mesh placed on the agar surface for haustorium induction assays. After overnight stratification at 4°C, agar plates were incubated at 25°C under long day conditions (16 hours light / 8 hours dark; LD). Horizontally-grown 6-day-old seedlings were used for transformation experiments. Vertically-grown 10-day-old seedlings were used for haustorium induction assays with chemical treatments. *A. thaliana* Col-0 wild type was used as the host plant in this study. *Arabidopsis* seeds were surface sterilized with 70% ethanol for 10 minutes and sown on half-strength MS agar plates (0.8% agar and 1% sucrose). After stratification at 4°C for 2 days, agar plates were incubated vertically at 22°C under LD conditions. Six-to eight-day-old seedlings were used for *P. japonicum* infection.

### Plasmid construction

Golden Gate cloning technology was used for plasmid construction (Engler et al., 2014). All *Bpi*I and *Bsa*I restriction sites were mutated in cloned DNA sequences. All primers used in this paper are listed in Supplementary table 1. Previously described golden gate modules were used to generate *DR5::3xmCherry-NLS*, *DR5::3xmCherry-NLS*+*PjCESA7::3xVenus-NLS*, *PjYUC3::3xVenus-NLS*, and *PjHB15a::3xVenus-SYP* (Ishida et al., 2016; Wakatake et al., 2018). *PjPIN1*, *PjPIN2*, *PjPIN3*, *PjPIN4*, *PjIPIN9*, *PjLAX1*, *PjLAX2*, *PjLAX4*, and *PjLAX5* promoter regions were PCR-amplified as multiple fragments from *P. japonicum* genomic DNA and cloned into pAGM1311. These amplified fragments were assembled into the pICH41295 level 0 vector. The *PjLAX3* promoter region was directly cloned into pICH41295 following PCR amplification from genomic DNA. The coding regions of *PIN* and *LAX* genes were also PCR-amplified as multiple fragments and inserted into pAGM1311. The Venus coding sequence was cloned into pAGM1311 using primers containing linker sequences (GGGGGA at the 5’ end and AGAAAAAGA at the 3’ end). Fragmented coding regions of *PIN* and *LAX* genes were integrated with the Venus coding sequence flanked by linker sequences into level 0 pICH41308 to generate fluorescent protein-fused sequences. The promoter sequence of pICH41295, the coding region with its fluorescent protein sequence in pICH41308, and the 3’UTR and terminator regions of pICH41276 were combined into level 1 vectors. These sequences were further subcloned into pAGM1311 to change antibiotic resistance markers for *Agrobacterium* selection. For RNAi experiments, we used pHG8-YFP for visual screening of hairy roots (Bandaranayake et al., 2010). Target sequences were PCR-amplified from *P. japonicum* genomic DNA and cloned into the pENTR vector (Thermo Fisher Scientific), then transferred into pHG8-YFP by the Gateway reaction using LR Clonase II Plus enzyme (Thermo Fisher Scientific).

### *P. japonicum* transformation

*P. japonicum* transformation was performed as previously described using *Agrobacterium rhizogenes* strain AR1193 (Ishida et al., 2011). Six-day-old *P. japonicum* seedlings were sonicated for 10 seconds, followed by vacuum infiltration for 5 minutes in an aqueous suspension of *A. rhizogenes*. Seedlings were then co-incubated with *A. rhizogenes* on Gamborg’s B5 medium (0.8% agar, 1% sucrose) containing 450 µM acetosyringone (Sigma-aldrich) at 22°C in the dark for 2 days, transferred to Gamborg’s B5 medium (0.8% agar, 1% sucrose) containing 300 µg/ml cefotaxime (Tokyo Chemical Industry), and incubated at 25°C under LD conditions.

### Confocal microscopy

Confocal microscopy was performed as previously described (Wakatake et al., 2018). *P. japonicum* plants with hairy roots were transferred onto 0.7% water agar plates and incubated at 25°C under LD conditions for 2 days. Freshly elongated hairy roots exhibiting fluorescence were placed in small glass petri dishes (IWAKI, Japan). Hairy roots were placed directly on the glass bottom of the dish and covered by a thin agar layer (0.7%). Plates were incubated overnight at 25°C under LD conditions. The following day, seven-day-old *Arabidopsis* seedlings were placed next to the hairy roots under the thin agar layers to induce haustorium formation, and then incubated in a growth chamber at 25°C under LD conditions. A Leica SP5 inverted confocal microscope was used for imaging. The Venus fluorescent protein was excited using a 514 nm laser, and emission was detected at 525-560 nm. The mCherry fluorescent protein was excited at 543 nm, and emission was detected at 570-640 nm. We selected hairy roots with strong fluorescence to clarify expression in deeper tissues for use in the figures shown in this study.

### Auxin transport inhibitor treatment

Ten-day-old *P. japonicum* seedlings grown on nylon mesh were transferred from half-strength MS medium to 0.7% agar plates together with underlying nylon mesh and incubated for 2 days at 25°C under LD conditions. *A. thaliana* seedlings were placed to next to *P. japonicum* seedlings on the nylon mesh so that *P. japonicum* root tips were in contact with *A. thaliana* roots. Plants were transferred simultaneously by placing the mesh onto agar media containing the auxin transport inhibitors 5 µM NPA or 10 µM CHAPAA (Sigma-Aldrich) at 0, 24, and 48 hpi. For local chemical treatments, small agar blocks containing 5 µM NPA were placed at the indicated sites shown in Fig. S1. Haustorium tissues were sampled at 96 hpi. Xylem bridge formation was examined as previously described (Wakatake et al., 2018).

### Phylogenetic analysis

Full-length amino acid sequences were aligned using MAFFT (v7.407) with the auto mode (Katoh and Standley, 2013). Alignment data were trimmed using trimAl (v1.4) (Capella-Gutierrez et al., 2009), then subjected to phylogenic analysis using IQ-TREE (v1.5.5) with default settings (Nguyen et al., 2015).

### Phenotyping and RT-qPCR for RNAi hairy roots

Freshly elongated YFP-positive hairy roots were transferred to small glass petri dishes, and parasitized *A. thaliana* roots were prepared as described in the confocal microscopy section above. Xylem bridge connections were examined under the confocal microscope at 96 hpi. Subsequently, haustorial tissues were dissected and pooled in a single tube to produce one biological replicate. Total RNA was extracted using a RNeasy Plant mini kit (Qiagen) from fine-powdered frozen tissue. cDNA was synthesized using ReverTraAce qPCR RT Master Mix (TOYOBO) and quantified with Thunderbird SYBR qPCR Mix (TOYOBO) using Stratagene mx3000p. *PjUBC2* was used as an internal control as previously described (Spallek et al., 2017).

## Accession number

Gene sequences are available under the GenBank accession numbers as follows: LC506019 (PjPIN1), LC5060120 (PjPIN2), LC506021 (PjPIN3), LC506022 (PjPIN4), LC506023 (PjPIN9), LC506024 (PjLAX1), LC506025 (PjLAX2), LC506026 (PjLAX3), LC506027 (PjLAX4), LC506028 (PjLAX5)

## Acknowledgements

We thank Max Fishman for critical reading of the manuscript.

## Author contributions

Conceptualization: T.W., S.Y., K.S.; Methodology: T.W., S.Y.; Software: T.W.; Validation: T.W.; Formal analysis: T.W.; Investigation: T.W.; Resources: T.W.; Data curation: T.W.; Writing - original draft: T.W., K.S.; Writing - review & editing: T.W., S.Y., K.S.; Visualization: T.W.; Supervision: S.Y., K.S.; Project administration: S.Y., K.S.; Funding acquisition: S.Y., K.S.

---

**Fig. S1.**
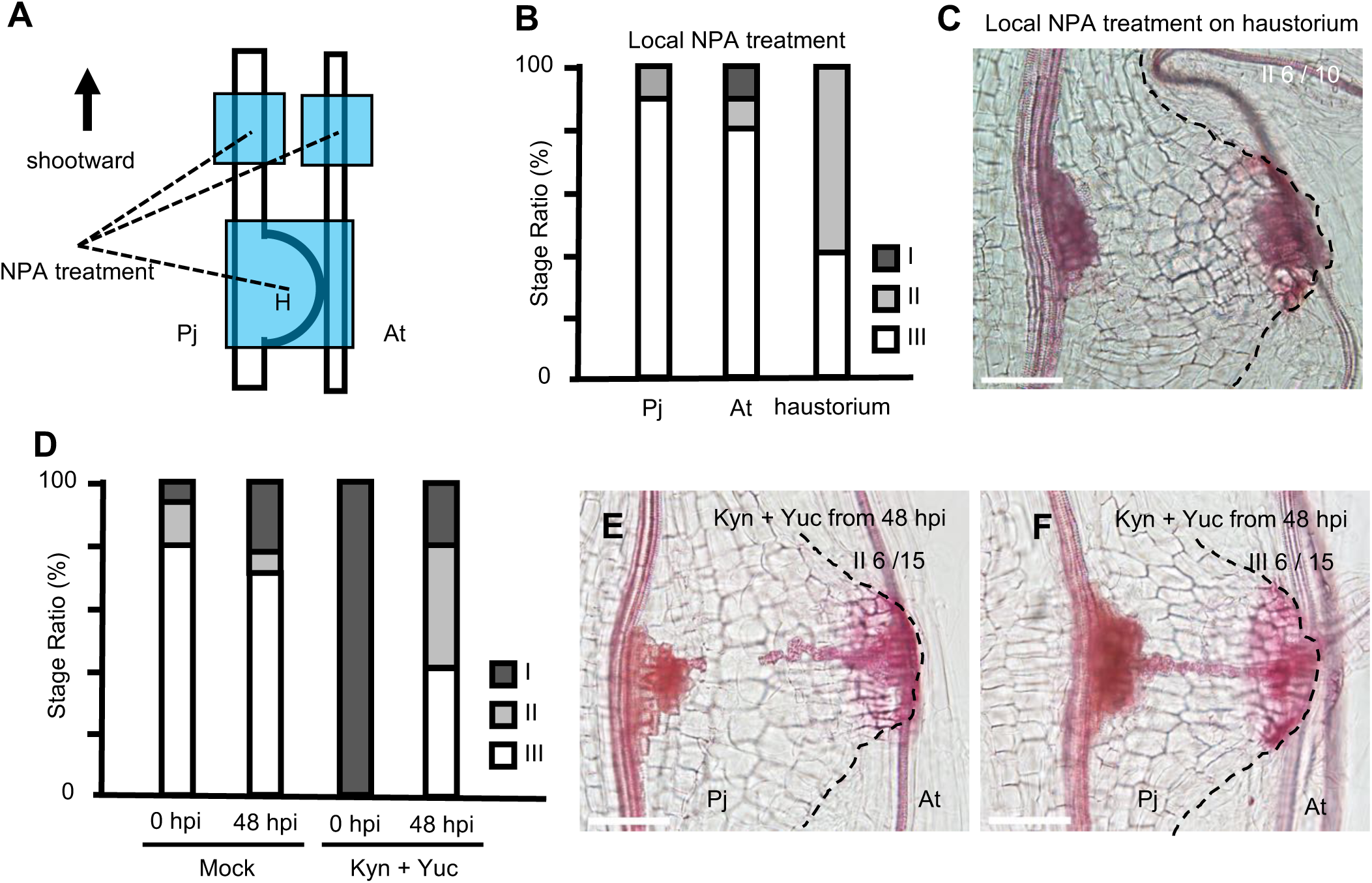
Local applications of an auxin transport inhibitor and auxin biosynthesis inhibitors. (A) The experimental design of local application of NPA (5 µM). H, haustorium. (B) Ratio of each developmental stage of xylem bridge formation at 96 hpi with local NPA treatments at 48 hours post infection (hpi) (I, pre-initiation; II, developing; III, fully connected). n = 10 ∼ 18 for each chemical treatment. (C) A representative photo of xylem bridge formation classified as stage II for local NPA treatment. (D) Ratio of each developmental stage of xylem bridge formation at 96 hpi with 10 µM L-Kynurenine (Kyn) + 20 µM Yuccasin (Yuc) treatments. n = 14-15 for each treatment. (E) and (F) Representative micrographs of xylem bridge development stages II (E) and III (F) for Kyn+Yuc treatment. White dotted lines mark the outline of haustoria. Tracheary elements are stained red with Safranin-O. Pj, *P. japonicum* root; At, *A. thaliana* root. Bar = 100 µm.

**Fig. S2.**
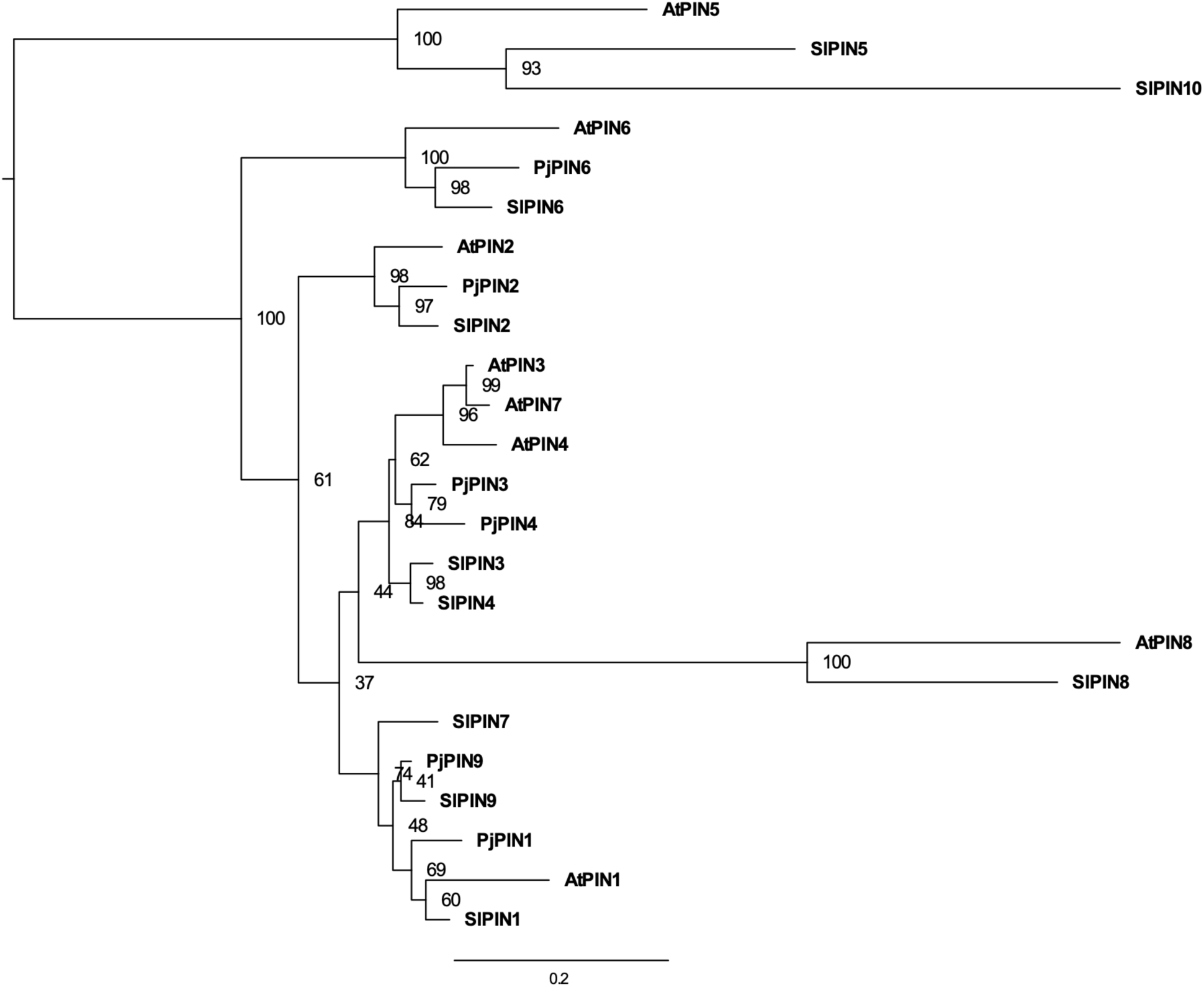
Phylogenetic tree for *A*. *thaliana* (At), *P*. *japonicum* (Pj), and *S*. *lycopersicum* (Sl) PIN family proteins. Full amino acid sequences of PIN proteins from three plant species, *A. thaliana* (At), *S. lycopersicum* (Sl), and *P. japonicum* (Pj), were used to construct a phylogenetic tree of PIN family proteins. Bootstrap values are indicated at the nodes. Amino acid substitutions per site are shown on the scale bar.

**Fig. S3.**
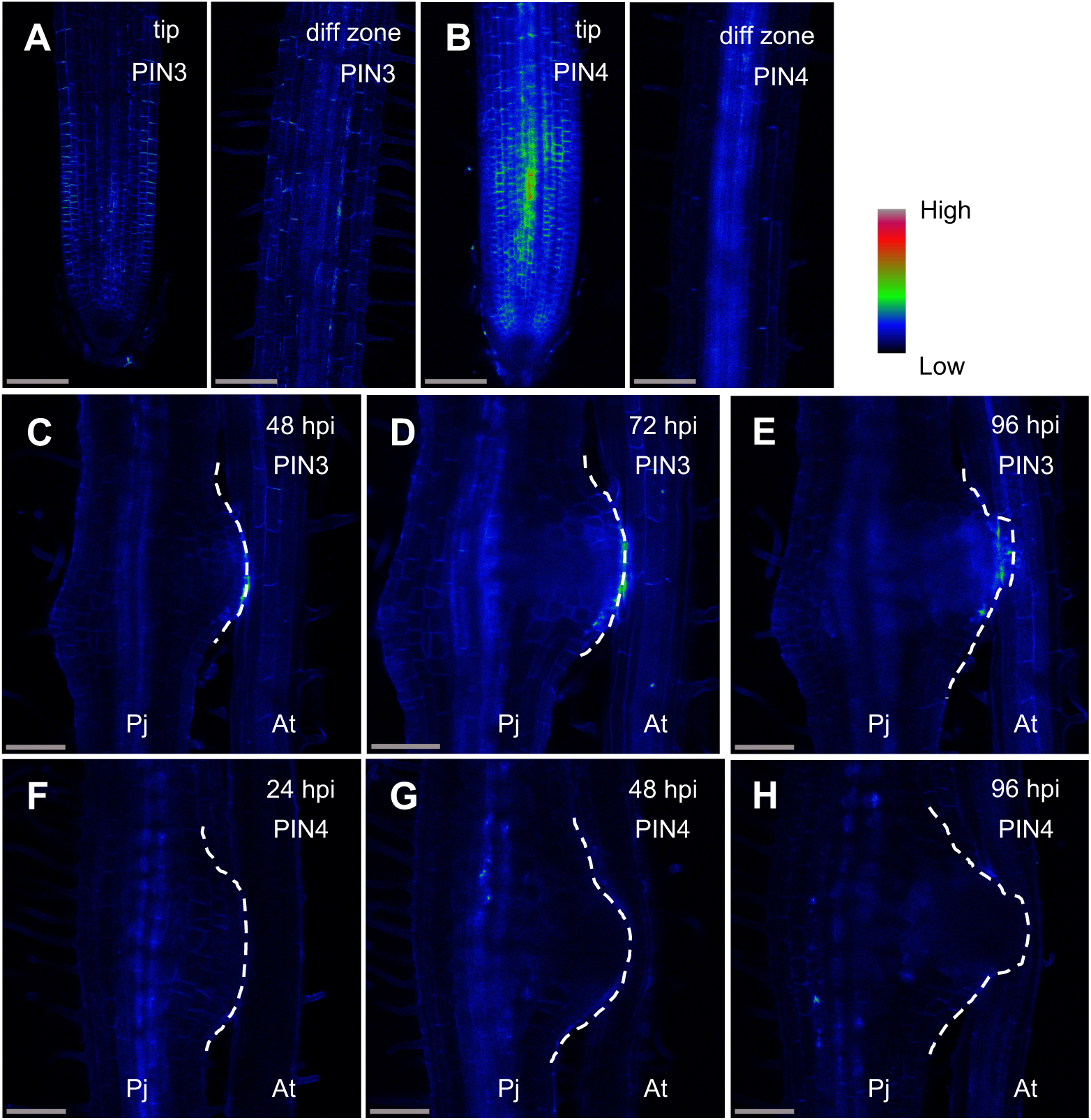
Expression patterns of PjPIN3 and PjPIN4 in *P*. *japonicum* root and haustoria. (A) and (C)-(E) *PjPIN3::PjPIN3-Venus* expression in *P. japonicum* root tips and differentiation zone (A), and during haustorium formation at 48, 72 and 96 hpi (C)-(E). (B) and (F)-(H) *PjPIN4::PjPIN4-Venus* expression in *P. japonicum* root tip (C) and during haustorium formation at indicated time points (F)-(H). Venus fluorescence intensity is depicted in a Rainbow RGB spectrum. The white dotted lines mark outlines of haustoria. hpi, hours post infection; Pj, *P. japonicum* root; At, *A. thaliana* root. Bar = 100 µm.

**Fig. S4.**
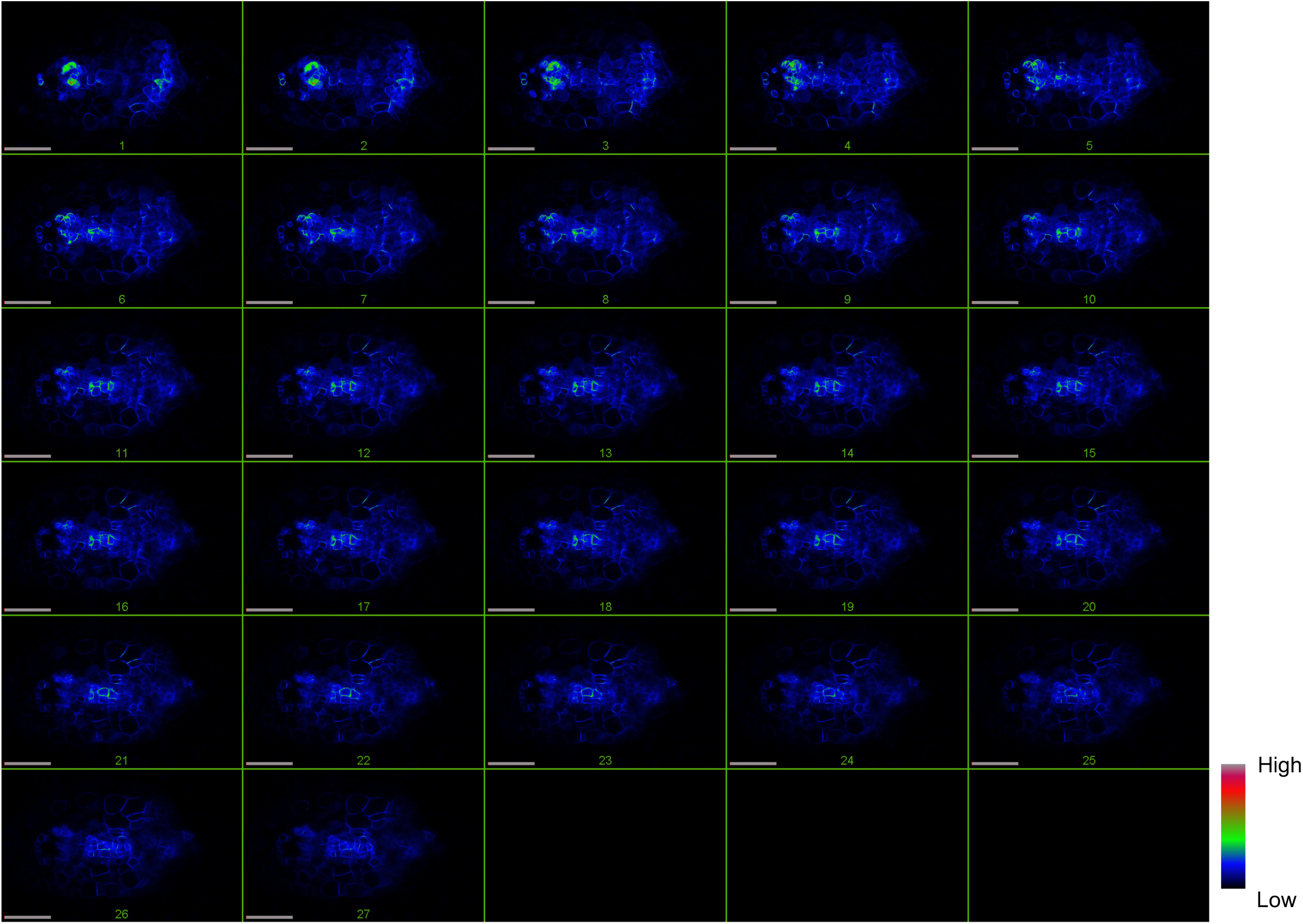
Subcellular localization of PjPIN1 in haustoria prior to xylem bridge formation. *PjPIN1::PjPIN1-Venus* expression in cross sections of haustoria at 72 hpi, but prior to xylem bridge formation. Venus fluorescence intensity is depicted in a Rainbow RGB spectrum. White lines mark outlines of haustoria. Pj, *P. japonicum* root; At, *A. thaliana* root. Bar = 100 µm.

**Fig. S5.**
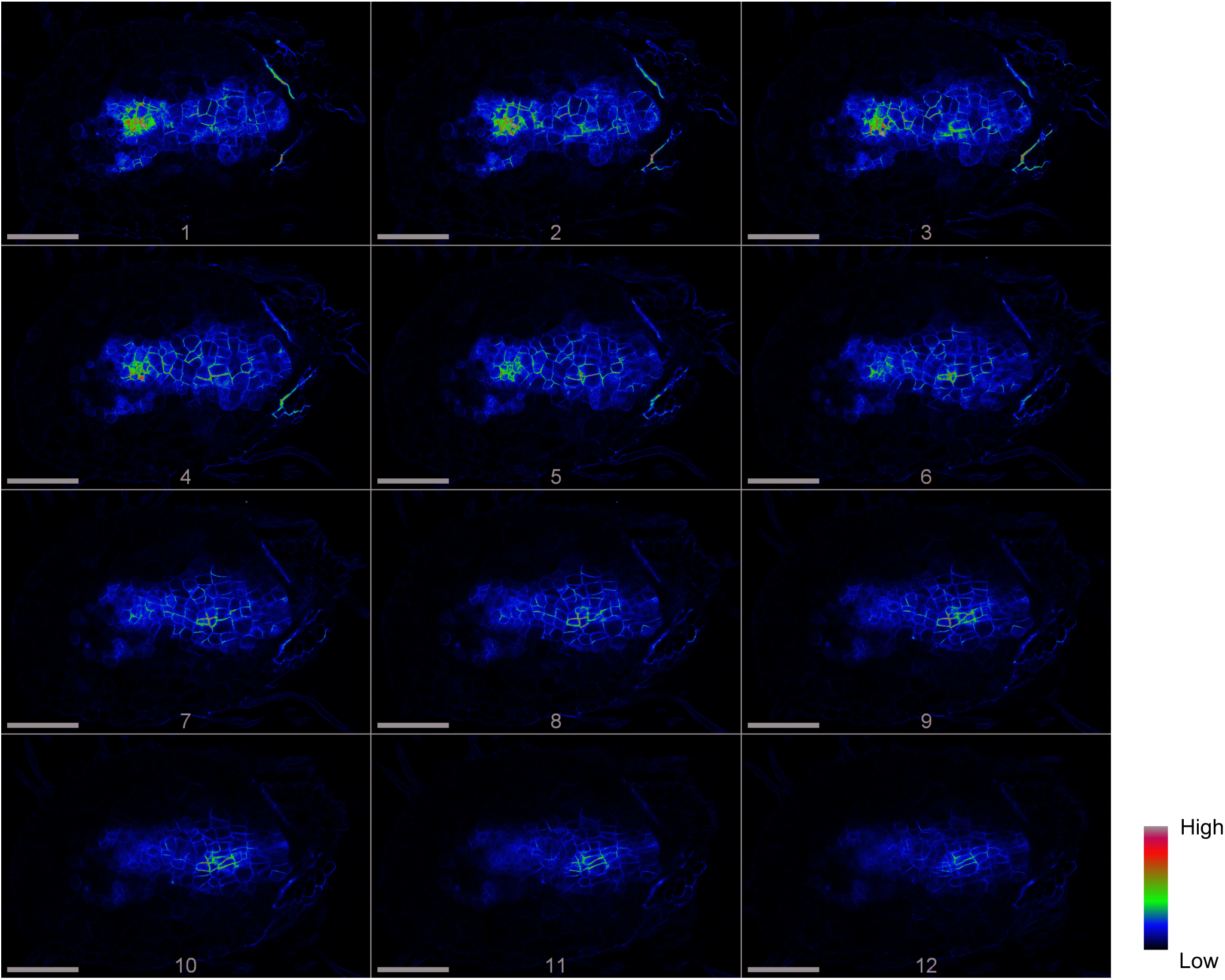
Subcellular localization of PjPIN9 in haustoria prior to xylem bridge formation. Full stack images are shown of *PjPIN9::PjPIN9-Venus* expression in haustorium cross sections at 72 hpi, but prior to xylem bridge formation. Venus fluorescence intensity is depicted in a Rainbow RGB spectrum. White lines mark outlines of haustoria. Pj, *P. japonicum* root; At, *A. thaliana* root. Bar = 100 µm.

**Fig. S6.**
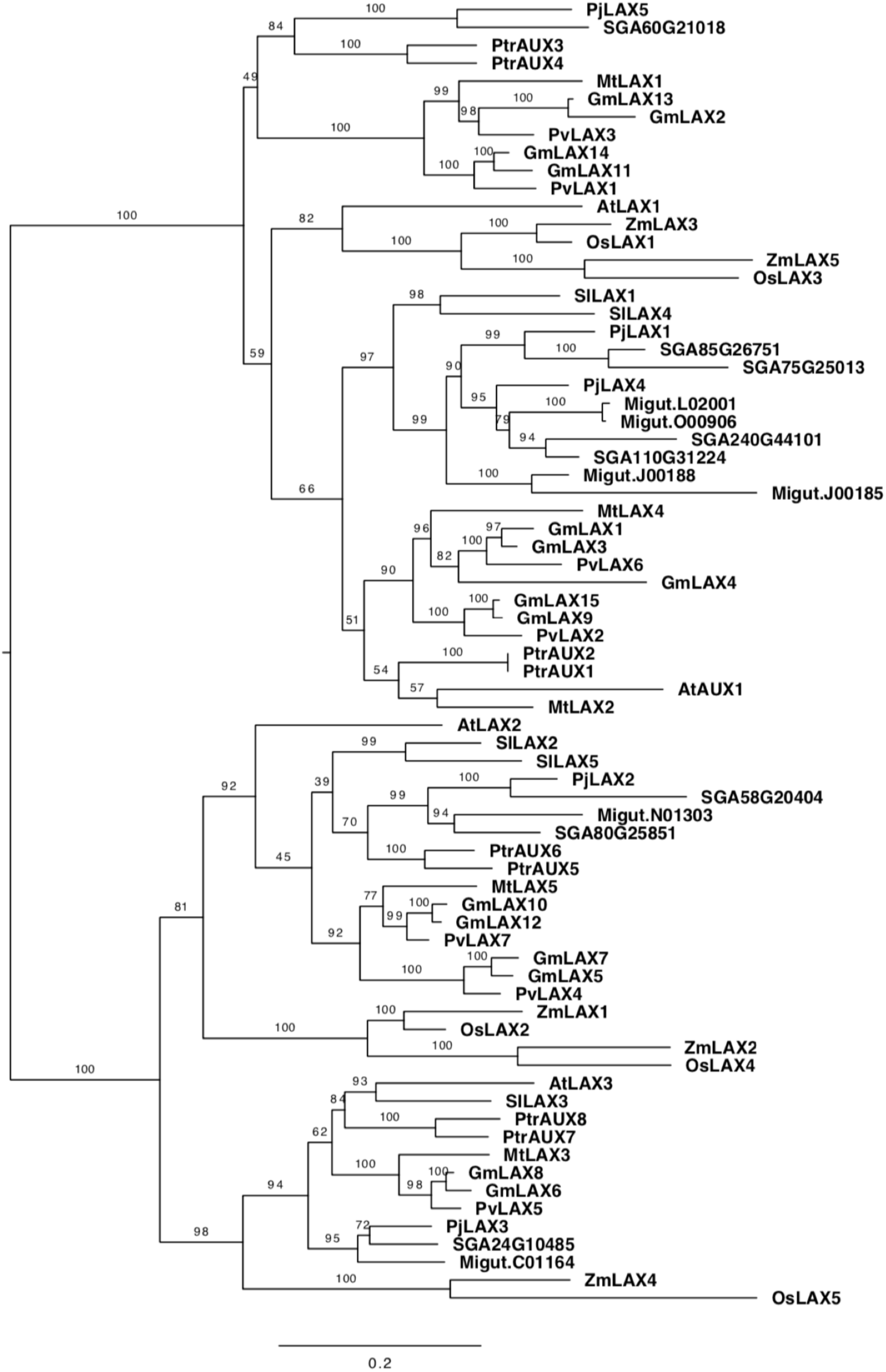
Phylogenetic tree of the AUX/LAX family. A. thaliana (At), *S. lycopersicum* (Sl), and *P. japonicum* (Pj), *Striga asiatica* (SGA), *Mimulus guttatus* (Migut), *Glycine max* (Gm), *Medicago truncatula* (Mt), *Phaseolus vulgaris* (Pv), *Populus trichocarpa* (Ptr), *Oryza sativa* (Os), *Zea maize* (Zm), were used to construct a phylogenetic tree of the AUX/LAX family. Bootstrap values are indicated at the nodes. Amino acid substitutions per site are shown on the scale bar.

**Fig. S7.**
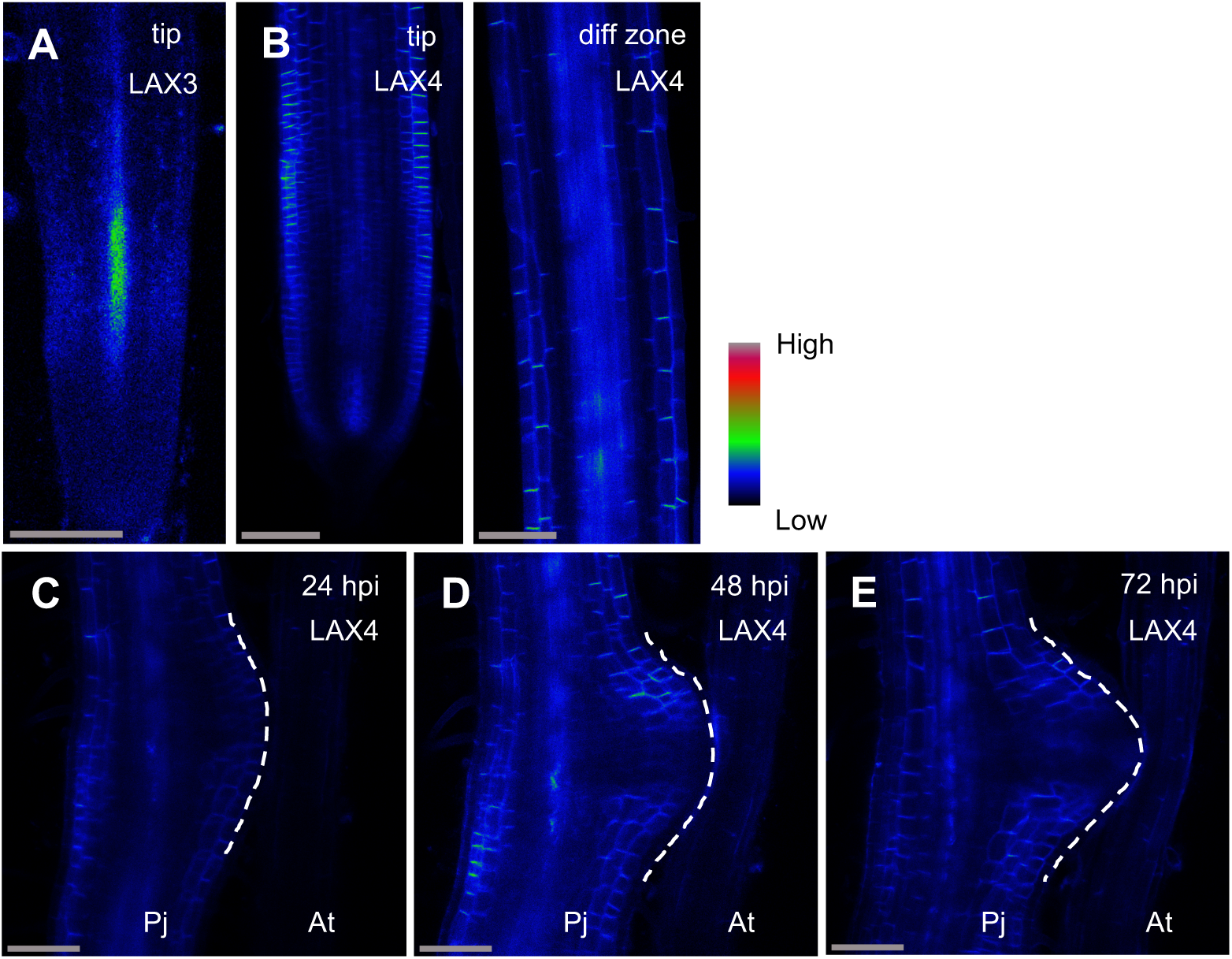
Expression patterns of PjLAX3 and PjLAX4 in *P*. *japonicum* roots and haustoria. (A) *PjLAX3::PjLAX3-Venus* expression in a *P. japonicum* root. PjLAX3 was expressed very weakly in vascular tissue of the basal meristem. However, no expression was detected in haustorium tissue. (B)-(E) *PjLAX4::PjLAX4-Venus* expression in *P. japonicum* root (B) and during haustorium formation at 24, 48 or 72 hpi (C)-(E). PjLAX4 was expressed in the epidermis, the outer cortex, and stele tissue, although no significant change was detected during haustorium formation. White dotted lines mark the outline of haustoria. Venus fluorescence is shown in Rainbow RGB spectrum. hpi, hours post infection; Pj, *P. japonicum* root; At, *A. thaliana* root. Bar = 100 µm.

**Fig. S8.**
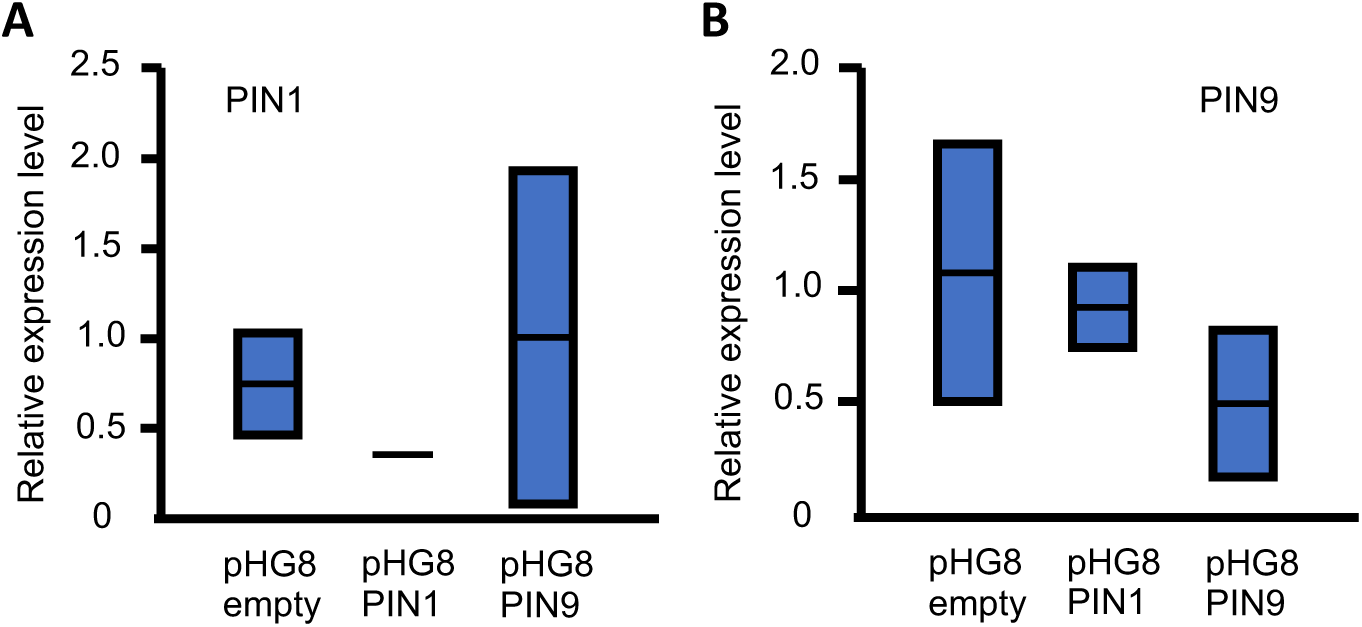
Reduced expression of PjPIN1 and PjPIN9 in knockdown roots. (A) and (B) Relative transcriptional levels of *PjPIN1* (A) and *PjPIN9* (B) in haustoria at 96 hpi. Means and values from two biological replicates are plotted. Each biological replicate includes 9-18 haustoria. *PjUBC2* transcription was used as an internal control.

**Supplemental Table 1.**
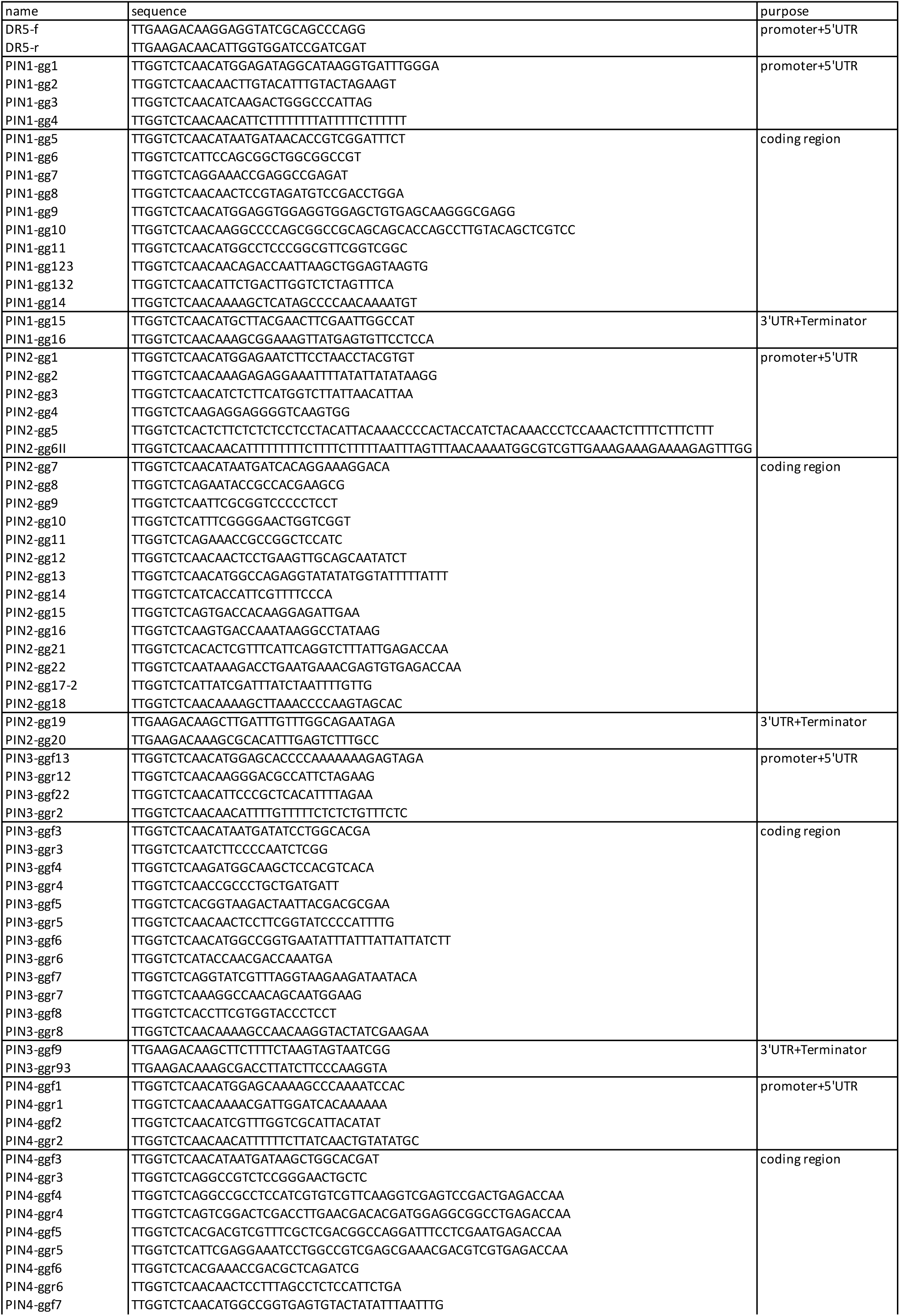
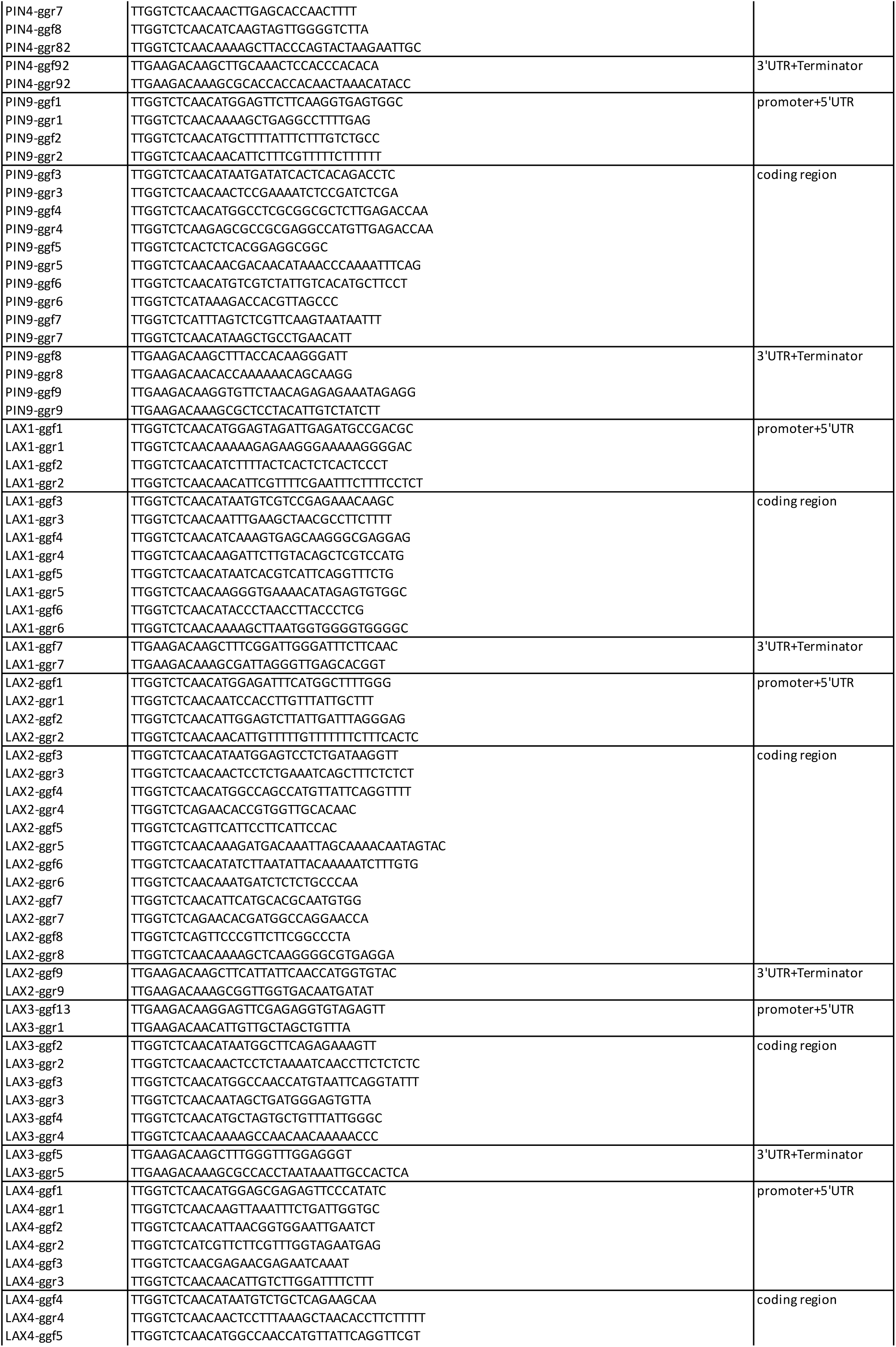
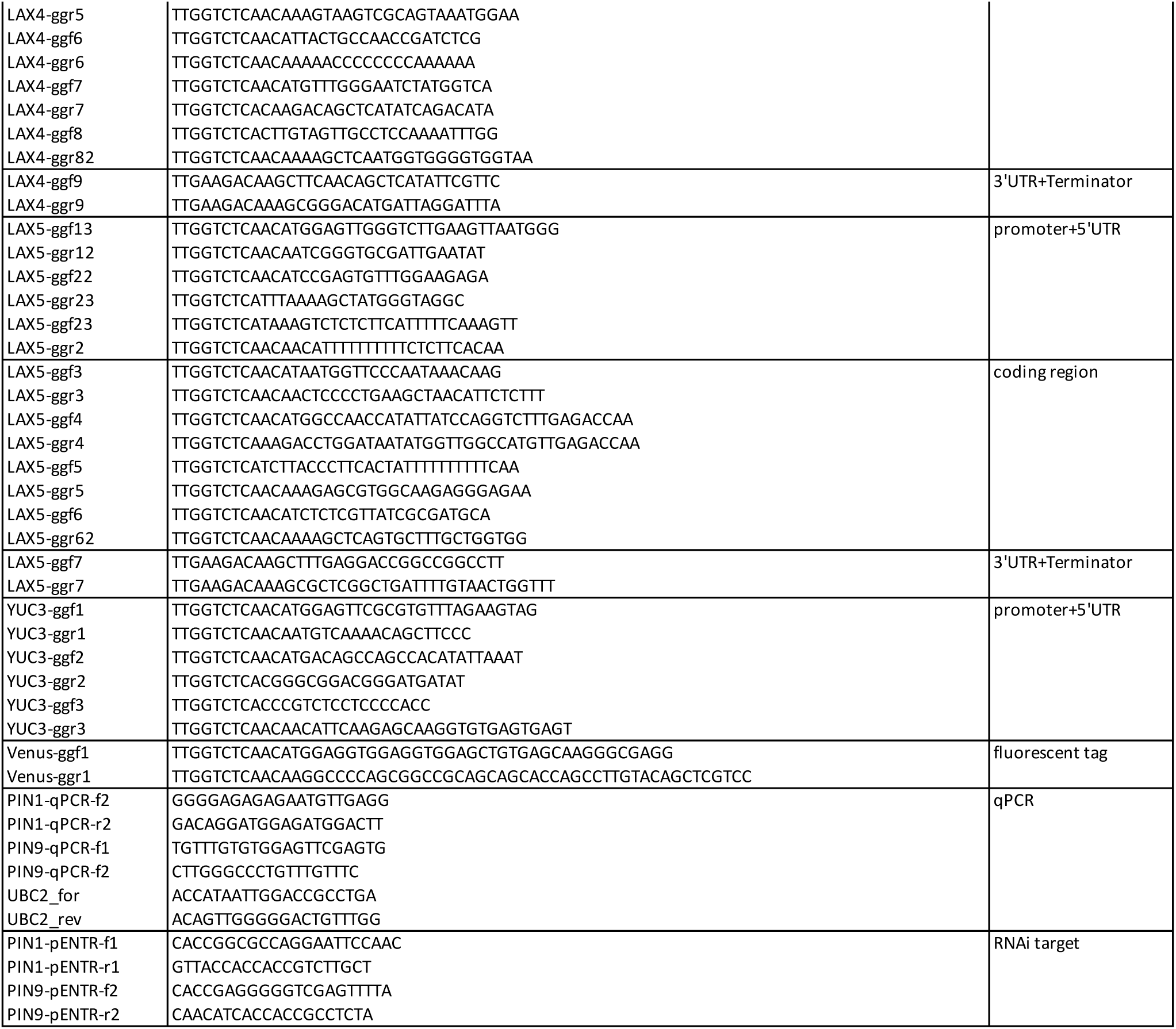
Primer list

